# Orai3 and Orai1 are essential for CRAC channel function and metabolic reprogramming in B cells

**DOI:** 10.1101/2022.05.06.490918

**Authors:** Scott M. Emrich, Ryan E. Yoast, Xuexin Zhang, Adam J. Fike, Yin-Hu Wang, Kristen N. Bricker, Anthony Tao, Ping Xin, Vonn Walter, Martin T. Johnson, Trayambak Pathak, Adam C. Straub, Stefan Feske, Ziaur S.M. Rahman, Mohamed Trebak

## Abstract

The essential role of store-operated Ca^2+^ entry (SOCE) through Ca^2+^ release-activated Ca^2+^ (CRAC) channels in T cells is well established. In contrast, the contribution of individual Orai isoforms to SOCE and their downstream signaling functions in B cells are poorly understood. Here, we demonstrate changes in expression of Orai isoforms in response to B cell activation. We show that Orai3 and Orai1 are essential components of native CRAC channels in B cells and are critical for primary B cell proliferation and survival. The combined loss of Orai1 and Orai3 strongly impairs SOCE, nuclear factor for activated T cells (NFAT) activation, mitochondrial respiration, glycolysis, and the metabolic reprogramming of B cells in response to antigenic stimulation. Our results clarify the molecular composition and cellular functions of SOCE in B lymphocytes.

## Introduction

Calcium (Ca^2+^) is an essential regulator of immune cell function. Crosslinking of immunoreceptors like the T cell receptor (TCR) or B cell receptor (BCR) triggers a robust elevation in intracellular Ca^2+^ concentrations through the release of endoplasmic reticulum (ER) Ca^2+^ stores and concomitant influx of Ca^2+^ from the extracellular space (Baba & Kurosaki, 2016; King & Freedman, 2009). In lymphocytes, Ca^2+^ entry across the plasma membrane (PM) is predominately achieved through Ca^2+^ release-activated Ca^2+^ (CRAC) channels, which constitute the ubiquitous store-operated Ca^2+^ entry pathway (SOCE) (Prakriya & Lewis, 2015; Trebak & Kinet, 2019; Trebak & Putney, 2017; Vaeth *et al*, 2020). Stimulation of immunoreceptors coupled to phospholipase C isoforms results in production of the secondary messenger inositol-1,4,5-trisphosphate (IP_3_), which triggers the release of ER Ca^2+^ through the activation of ER-resident IP_3_ receptors. A reduction in ER Ca^2+^ concentrations is sensed by stromal interaction molecule 1 (STIM1) and its homolog STIM2, resulting in a conformational change and clustering in ER-PM junctions where STIM molecules interact with and activate PM hexameric Orai channels (Orai1-3) to mediate SOCE(Lunz *et al*, 2019). Ca^2+^ entry through CRAC channels regulates immune cell function through a host of Ca^2+^-sensitive transcription factors including nuclear factor of activated T cells (NFAT), nuclear factor κB (NF-κB), c-Myc, and mTORC1(Berry *et al*, 2018; Trebak & Kinet, 2019; Vaeth & Feske, 2018; Vaeth *et al*., 2020; Vaeth *et al*, 2017a). The coordination of these master transcriptional regulators is indispensable for innate and adaptive immune cell effector function including entry into the cell cycle, clonal expansion, cytokine secretion, differentiation, and antibody production(Shaw & Feske, 2012; Trebak & Kinet, 2019; Vaeth *et al*., 2020).

Orai1 has long been firmly established as a central component of the *native* CRAC channel in all cell types studied. Although all three Orai isoforms are ubiquitously expressed across tissue types and can form functional CRAC channels when ectopically expressed, only recently Orai2 and Orai3 have emerged as critical regulators of native CRAC channel function (Emrich *et al*, 2022; Tsvilovskyy *et al*, 2018; Vaeth *et al*, 2017b; Yoast *et al*, 2020a). Patients with inherited loss-of-function (LoF) mutations in *Orai1* (e.g. R91W mutation) develop a CRAC channelopathy with symptoms including combined immunodeficiency, ectodermal dysplasia, muscular hypotonia, and autoimmunity(Feske *et al*, 2006; Lian *et al*, 2017; McCarl *et al*, 2009). While these patients display relatively normal frequencies of most immune cell populations, they are highly susceptible to reoccurring viral, bacterial, and fungal infections due to defects in T cell expansion, cytokine secretion, and metabolism(Vaeth *et al*., 2020). There are currently no reported patient mutations in *Orai2* or *Orai3* genes and the role these channels play within the immune system had largely been unclear. Interestingly, recent studies utilizing global Orai2 knockout mice have demonstrated that loss of Orai2 leads to enhanced SOCE and corresponding CRAC currents in bone marrow-derived macrophages, dendritic cells, T cells, enamel cells, and mast cells(Eckstein *et al*, 2019; Tsvilovskyy *et al*., 2018; Vaeth *et al*., 2017b), suggesting that Orai2 is a negative regulator of CRAC channel activity. Combined knockout in mice of both Orai1 and Orai2 in T cells led to a near ablation of SOCE and impaired humoral immunity, while ectopic expression of pore-dead mutants of Orai1 (E106Q) or Orai2 (E80Q) into individual Orai1^-/-^ or Orai2^-/-^ T cells blocked native SOCE(Vaeth *et al*., 2017b). Similarly, generation of HEK293 cell lines lacking each Orai isoform individually and in combination resulted in altered Ca^2+^ oscillation profiles, CRAC currents, and NFAT isoform activation (Emrich *et al*., 2022; Emrich *et al*, 2021; Yoast *et al*., 2020a; Yoast *et al*, 2020b). These data provide strong evidence that Orai2 and Orai3 exert negative regulatory effects on native CRAC channels likely by forming heteromeric CRAC channels with Orai1.

In contrast to T cells, much less is known regarding the role of SOCE in B lymphocytes. Early landmark work investigating B cell-specific STIM knockout mice established that STIM1 mediates the vast majority of SOCE in B cells, while only the combined deletion of both STIM1 and STIM2 resulted in substantial impairments in B cell survival, proliferation, and NFAT-dependent IL-10 secretion (Matsumoto *et al*, 2011). Unexpectedly, mice with STIM1/STIM2-deficient B cells show relatively normal humoral immune responses to both T cell-dependent and independent immunizing antigens. These and other studies suggest that the severe immunodeficiency observed in CRAC channelopathy patients is due to impaired T cell responses (Matsumoto *et al*., 2011; Vaeth *et al*, 2016). Investigation of Orai1^-/-^ mice on the mixed Institute for Cancer Research (ICR) background and Orai1^R93W^ knock-in mice (the equivalent of human R91W mutation) demonstrated that SOCE is significantly attenuated in B cells(Gwack *et al*, 2008; McCarl *et al*, 2010). However, SOCE in B cells is only partially reduced with the loss of Orai1, suggesting that Orai2 and/or Orai3 mediate the remaining SOCE in B cell populations. Thus, the contribution of each Orai isoform to native CRAC channel function and downstream signaling in B cells has remained obscure.

In this study, we investigated the contributions of Orai isoforms to SOCE and its downstream signaling in B cells through multiple CRISPR/Cas9 knockout B cell lines and novel B cell specific Orai knockout mice. Our findings demonstrate that expression of each Orai isoform is dynamically regulated in response to B cell activation and that the magnitude of SOCE is unique among effector B cell populations. We show that deletion of Orai1 alone does not alter BCR-evoked cytosolic Ca^2+^ oscillations, proliferation, and development. Unexpectedly, we reveal that Orai3 is an essential component of the native CRAC channel that coordinates with Orai1 to mediate the majority of SOCE in B cells, which is critical for B cell mitochondrial metabolism. Transcriptome and metabolomic analysis uncovered key signaling pathways that are regulated by SOCE and the phosphatase calcineurin for the efficient transition of B cells from a quiescent to metabolically active state. Our data elucidate the role of SOCE in B cell functions and show that CRAC channel function in B lymphocytes is mediated by both Orai1 and Orai3. This knowledge is crucial for potential targeting of CRAC channels in specific lymphocyte subsets in immune and inflammatory disease.

## Results

### Orai channel isoforms are dynamically regulated in response to B cell activation

Throughout their lifespan, B lymphocytes must undergo dramatic periods of metabolic adaptation whereby naïve, metabolically quiescent cells prepare to clonally expand and differentiate into effector populations (Akkaya & Pierce, 2019; Boothby & Rickert, 2017). Induction of these diverse metabolic programs is driven by downstream BCR-mediated Ca^2+^ signals in combination with Ca^2+^-independent costimulatory signals including CD40 activation and/or stimulation with various toll-like receptor (TLR) ligands (Akkaya *et al*, 2018; Baba & Kurosaki, 2016; Berry *et al*, 2020). Thus, using Seahorse assays we first evaluated how activation of wild-type mouse primary splenic B cells (isolated as described in methods) regulates oxygen consumption rates (OCR) and extracellular acidification rates (ECAR) as indicators of oxidative phosphorylation (OXPHOS) and glycolysis, respectively (**Fig. 1A, B**). Naïve primary B cells from mouse spleens were isolated by negative selection and stimulated for 24 hours under five different conditions as follows: 1) control unstimulated cells (Unstim), 2) stimulation with anti-IgM antibodies (IgM) to activate the BCR, 3) stimulation with anti-CD40 (CD40), 4) stimulation with lipopolysaccharides (LPS), and 5) co-stimulation with anti-IgM and anti-CD40. B cell stimulation for 24 hours led to a robust increase in basal OCR, except for B cells stimulated with anti-CD40 alone where the increase in basal OCR was not statistically significant (**Fig. 1A, C**). **Figure 1B** shows that B cells stimulated with either anti-IgM, anti-IgM + anti-CD40 or LPS become highly energetic and upregulate both OXPHOS and glycolytic pathways. However, B cells stimulated with anti-CD40 alone remain metabolically quiescent like control non-stimulated B cells (**Fig 1B**). We used the protonophore trifluoromethoxy carbonylcyanide phenylhydrazone (FCCP) to dissipate the mitochondrial membrane potential and calculate the maximal respiratory capacity of B cells. Consistent with results obtained for basal OCR, stimulated B cells showed robust increase in maximal respiratory capacity except for cells stimulated with anti-CD40 alone (**Fig. 1D**). Both total mitochondrial content and membrane potential measured with MitoTracker Green and TMRE staining respectively, were substantially increased following anti-IgM, anti-IgM + anti-CD40, or LPS stimulation while both parameters were only slightly increased with stimulation with anti-CD40 alone (**Fig. 1E, F**). While previous studies suggested that Orai1 regulates the majority of SOCE in primary B cells (Gwack *et al*., 2008; McCarl *et al*., 2010), the expression of different Orai isoforms both at rest and upon B cell activation is unknown. Utilizing the same five experimental conditions described above, we analyzed the mRNA expression of *Orai1, Orai2*, and *Orai3* following 24 hours of stimulation. Stimulation with either anti-IgM or anti-IgM + anti-CD40 dramatically increased *Orai1* expression, while the magnitude of this change following anti-CD40 or LPS stimulation was smaller and did not reach statistical significance (**Fig. 1G**). Interestingly, stimulation conditions that strongly increased mitochondrial respiration (anti-IgM, anti-IgM + anti-CD40, LPS) all significantly reduced expression of *Orai2* while stimulation with anti-CD40 alone had no significant effect on *Orai2* expression (**Fig. 1H**). The expression of Orai3 was significantly increased only with anti-IgM + anti-CD40 or LPS stimulation, but not with anti-IgM or anti-CD40 alone (**Fig. 1I**).

**Figure 1.**
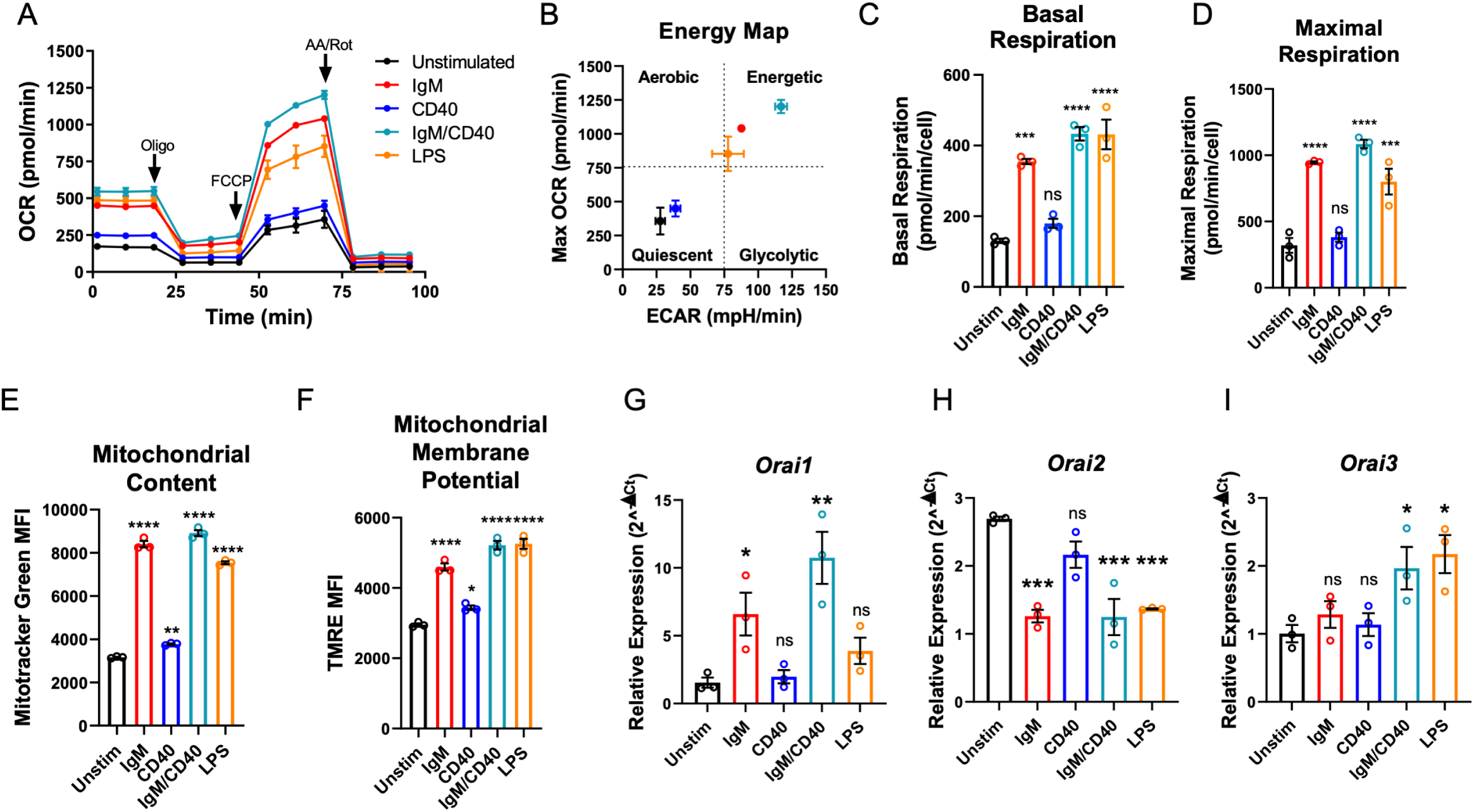
B cell activation dynamically regulates Orai channel expression. (A) Measurement of oxygen consumption rate (OCR) in primary B lymphocytes following 24-hour stimulation with anti-IgM (20 μg/mL), anti-CD40 (10μg/mL), anti-IgM + anti-CD40, or LPS (10μg/mL) using the Seahorse Mito Stress Test (n = 3 biological replicates). (B) Energy map of maximal OCR and extracellular acidification rate (ECAR) following addition of the protonophore carbonylcyanide p-trifluoromethoxyphenyl hydrazone (FCCP). (C, D) Quantification of basal (C) and maximal (D) respiration from Seahorse traces in (A) (One-way ANOVA with multiple comparisons to Unstimulated). (E, F) Measurement of (E) total mitochondrial content with the fluorescent dye MitoTracker Green and (F) mitochondrial membrane potential with TMRE following 24-hour stimulation (n = 3 biological replicates; One-way ANOVA with multiple comparisons to Unstimulated). (G-I) Quantitative RT-PCR of (G) *Orai1*, (H) *Orai2*, and (I) *Orai3* mRNA following 24-hour of stimulation with the stimuli indicated (n = 3 biological replicates; one-way ANOVA with multiple comparisons to Unstimulated). All scatter plots and Seahorse traces are presented as mean ± SEM. For all figures, *p<0.05; **p<0.01; ***p<0.001; ****p<0.0001; ns, not significant.

### Both Orai3 and Orai1 are essential components of SOCE in A20 B cells

Robust activation of primary B cells with anti-IgM/anti-CD40 co-stimulation resulted in upregulation of both *Orai1* and *Orai3* mRNA with strong downregulation of *Orai2* (**Fig. 1G-I**). We therefore reasoned that B cells may sustain long-term cytosolic Ca^2+^ signals through Orai1 and/or Orai3, while downregulation of Orai2 might relieve the inhibition of native CRAC channels, as recently demonstrated for activated T cells (Vaeth *et al*., 2017b). To gain insights into the molecular composition of CRAC channels in B cells, we first developed an *in vitro* system utilizing the mouse A20 B lymphoblast cell line to generate single and double Orai1 and Orai3 knockout cell lines with CRISPR/Cas9 technology. Two guide RNA (gRNA) sequences were utilized to cut at the beginning and end of the coding region of mouse *Orai1* and *Orai3* (*mOrai1* and *mOrai3*), effectively excising the entirety of the genomic DNA for each *Orai* gene (**Fig. 2A**). We generated multiple A20 clones that were lacking Orai1 and Orai3 individually and in combination and these knockout clones were validated through genomic DNA sequencing and the absence of *Orai1* or *Orai3* transcripts by qPCR (**Fig. 2B-D**). We measured SOCE in A20 cells after passive store depletion with thapsigargin, an inhibitor of the sarcoplasmic/endoplasmic reticulum ATPase (SERCA). A20 cells lacking Orai1 demonstrated a large reduction in maximal SOCE by ∼ 62%, while in cells lacking Orai3 SOCE was reduced by ∼28% (**Fig. 2E, F**). Importantly, the combined deletion of both Orai1 and Orai3 caused a near abrogation of SOCE by ∼ 91%. (**Fig. 2E, F**). Furthermore, measurements of cytosolic Ca^2+^ oscillations induced by anti-IgG stimulation demonstrated that the oscillation frequency was substantially reduced with combined Orai1/Orai3 knockout, while only slightly reduced with loss of either Orai1 or Orai3 individually (**Fig. 2G-K**). These data suggested that both Orai3 and Orai1 were crucial for optimal CRAC channel function in A20 B lymphocytes. These data are consistent with the function of Orai channels in other cell types. Indeed, we recently utilized a series of single, double, and triple Orai CRISPR/Cas9 knockout HEK293 cell lines and showed that SOCE, but not Orai1, is required for agonist induced Ca^2+^ oscillations (Yoast *et al*., 2020b). Under conditions of physiological agonist stimulation that causes modest ER store depletion while eliciting Ca^2+^ oscillations, Orai2 and Orai3 were sufficient to maintain cytosolic Ca^2+^ oscillations, while having relatively minor contributions to SOCE induced by maximal store depletion (Yoast *et al*., 2020b).

**Figure 2.**
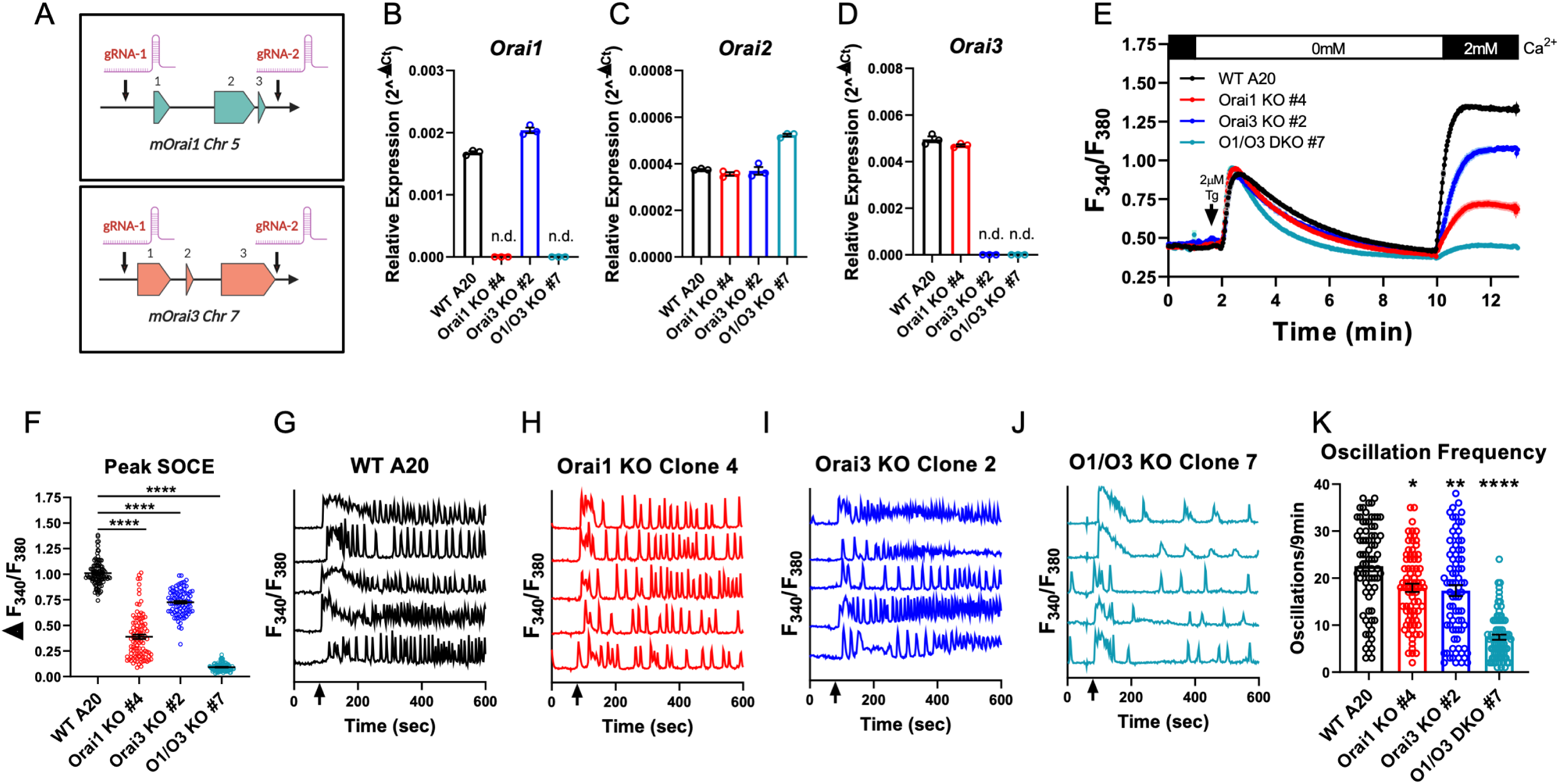
Orai1 and Orai3 mediate the bulk of SOCE in A20 B lymphoblasts. (A) Cartoon schematic of the two gRNA CRISPR strategy we used to excise mouse *Orai1* and *Orai3* genes. (B-D) Quantitative RT-PCR of (B) *Orai1*, (C) *Orai2*, and (D) *Orai3* mRNA in A20 Orai CRISPR clones (n = 3 biological replicates). (E) Measurement of SOCE with Fura2 upon store depletion with 2µM thapsigargin in 0mM Ca^2+^ followed by re-addition of 2mM Ca^2+^ to the external bath solution. (F) Quantification of peak SOCE in (E) (from left to right n = 99, 100, 89, and 98 cells; Kruskal-Wallis test with multiple comparisons to WT A20). (G-J) Representative Ca^2+^ oscillation traces from 5 cells/condition measured with Fura2 upon stimulation with 10 μg/mL anti-IgG antibodies at 60 seconds (indicated by arrows) in the presence of 2mM external Ca^2+^. (K) Quantification of total oscillations in 9 minutes from (G-J) (from left to right n = 76, 79, 79, and 78 cells; Kruskal-Wallis test with multiple comparisons to WT A20). All scatter plots are presented as mean ± SEM. For all figures, *p<0.05; **p<0.01; ****p<0.0001; ns, not significant.

### Orai1 is dispensable for cytosolic Ca^2+^ oscillations in primary B cells

To determine the contribution of Orai1-mediated Ca^2+^ signals to primary B cell function, we generated B cell specific Orai1 knockout (*Orai1*^*fl/fl*^ *Mb1*^*cre/+*^) mice (**Fig. 3**). Compared to *Mb1*^*cre/+*^ controls, the average surface expression of Orai1 was significantly reduced on B220^+^ B cells from *Orai1*^*fl/fl*^ *Mb1*^*cre/+*^ mice (**Fig. 3A, B**). Further, B cells from *Orai1*^*fl/fl*^ *Mb1*^*cre/+*^ mice showed a near abrogation of *Orai1* mRNA compared to *Mb1*^*cre/+*^ control mice (**Fig. 3C**) without any significant change in *Orai2* (**Fig. 3D**) or *Orai3* (**Fig. 3E**) mRNA expression. In *Mb1*^*cre/+*^ control mice, Orai1 expression was comparable between B220^+^ B cells and CD8^+^ T cells, with CD4^+^ T cells showing a slightly lower signal (**Fig. Fig. 3-Fig. Supplement A, B**). After 24-hour stimulation of B cells under the five conditions described above, Orai1-deficient B cells showed no apparent defects in activation as expression of MHC-II was comparable to control B cells (**Fig. 3-Fig. Supplement 1C-E**), while expression of CD86 was slightly reduced with anti-IgM stimulation or anti-IgM + anti-CD40 co-stimulation (**Fig. 3-Fig. Supplement 1F-H**). Measurement of SOCE demonstrated that Ca^2+^ entry was reduced by ∼69% in B cells isolated from *Orai1*^*fl/fl*^ *Mb1*^*cre/+*^ mice (**Fig. 3F, G**). We measured cytosolic Ca^2+^ oscillations in response to anti-IgM stimulation in the presence of 1 mM extracellular Ca^2+^. In agreement with previous reports and consistent with our data with A20 B lymphoblasts (**Fig. 2G, H**), Orai1-deficient B cells display a comparable frequency of agonist-induced Ca^2+^ oscillations as control B cells (**Fig. 3H-J**). Thus, while Orai1 knockout significantly decreased SOCE in primary B cells, it is dispensable for maintaining Ca^2+^ oscillations in response to physiological agonist stimulation.

**Figure 3.**
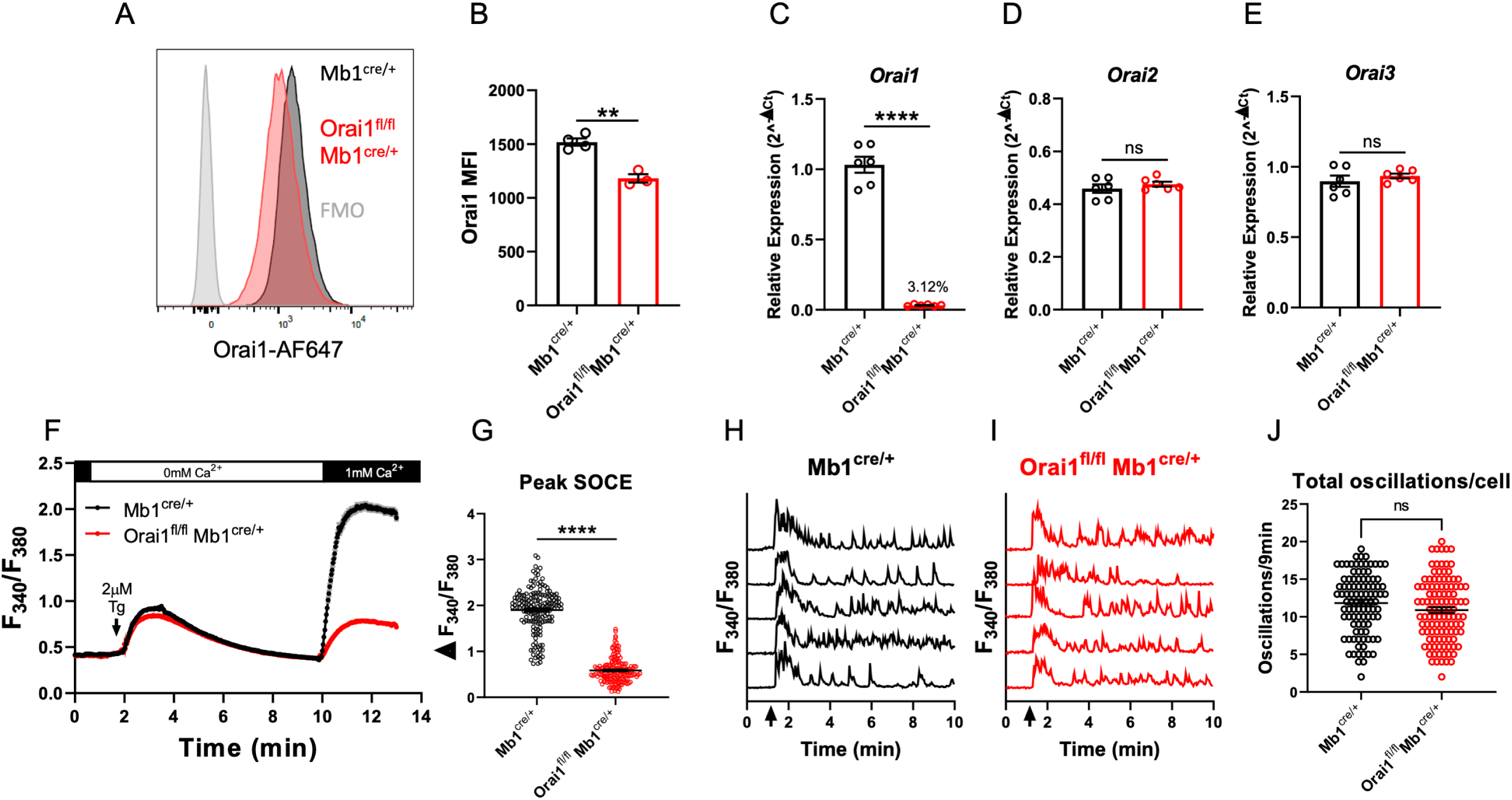
Orai1 is dispensable for BCR-induced Ca^2+^ oscillations in primary B cells. (A) Representative FACS plot of B cells isolated from *Mb1*^*cre/+*^ and *Orai1*^*fl/fl*^ *Mb1*^*cre/+*^ mice and stained with anti-Orai1 antibodies. (B) Quantification of Orai1 mean fluorescence intensity (MFI) on B cells from *Mb1*^*cre/+*^ and *Orai1*^*fl/fl*^ *Mb1*^*cre/+*^ mice shown in (A) (n = 4 and 3 biological replicates; unpaired T-test). (C-E) Quantitative RT-PCR of (C) *Orai1*, (D) *Orai2*, and (E) *Orai3* mRNA in isolated B cells (n = 6 biological replicates for each; Mann-Whitney test). (F) Measurement of SOCE with Fura2 upon store depletion with 2µM thapsigargin in 0mM Ca^2+^ followed by re-addition of 1mM Ca^2+^ to the external bath solution. (G) Quantification of peak SOCE in (F) (n = 169 and 178 cells; Mann-Whitney test. (H-I) Representative Ca^2+^ oscillation traces from 5 cells/condition measured with Fura2 upon stimulation with 20 μg/mL anti-IgM antibodies at 1 minute (indicated by arrows) in the presence of 1mM external Ca^2+^. (J) Quantification of total oscillations in 9 minutes from (I, J) (n = 97 and 111 cells; Mann-Whitney test). All scatter plots are presented as mean ± SEM. For all figures, **p<0.01; ****p<0.0001; ns, not significant.

### Loss of Orai1 does not alter B cell development

While patients and mouse models with loss of function (LoF) mutations in *Orai1* present with severe immunodeficiency due to impaired T cell function, the development of most immune cell populations is largely unaltered, suggesting SOCE is dispensable for initial immune cell selection and development (Gwack *et al*., 2008; Lacruz & Feske, 2015; McCarl *et al*., 2010). We evaluated B cell development within the bone marrow and spleen of *Orai1*^*fl/fl*^ *Mb1*^*cre/+*^ mice (**Fig. 4**). In agreement with previous reports that investigated B cell development in global *Orai1* deficient mice, development of early B cell progenitors in fractions A-F within the bone marrow were unaltered in *Orai1*^*fl/fl*^ *Mb1*^*cre/+*^ mice (**Fig. 4A-C**). Similarly, we observed no significant differences in peripheral transitional type 1 (T1), T2, and T3 immature B cells in the spleen (**Fig. 4D, F**). Mature B cell populations in the spleen are predominately follicular B cells with a smaller fraction of marginal zone B cells, and these ratios were largely comparable between control and *Orai1*^*fl/fl*^ *Mb1*^*cre/+*^ mice (**Fig. 4E, F**).

**Figure 4.**
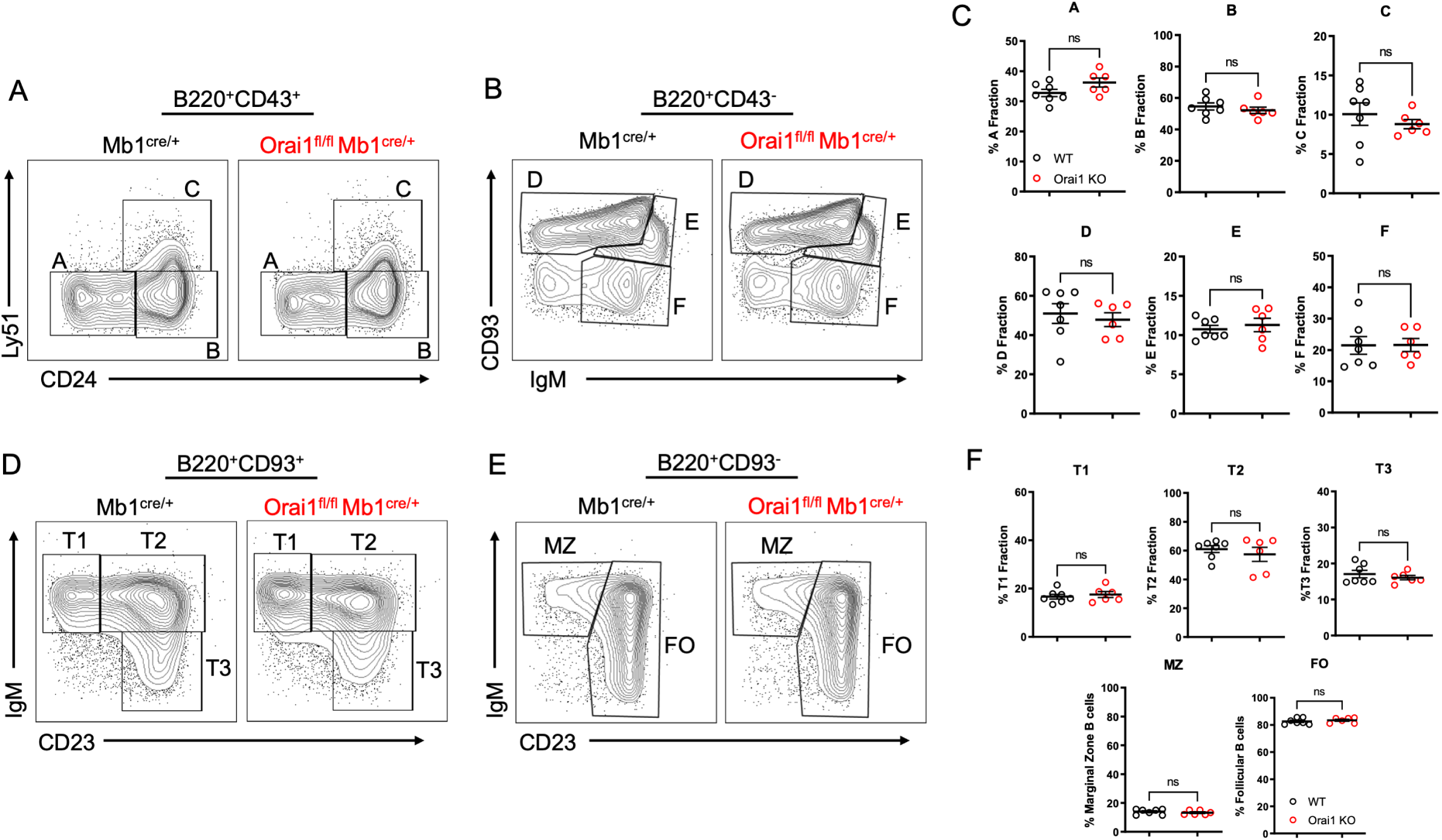
Orai1 is dispensable for B cell development. (A-B) Flow cytometric analysis of bone marrow populations for B cell fractions A (B220^+^CD43^+^HSA^-^BP-1^-^), B (B220^+^CD43^+^HSA^+^BP-1^-^), C (B220^+^CD43^+^HSA^+^BP-1^+^), D (B220^+^CD43^-^IgM^-^CD93^+^), E (B220^+^CD43^-^IgM^+^CD93^+^), and F (B220^+^CD43^-^IgM^+^CD93^-^). (C) Quantification of bone marrow populations in (A, B) (n = 7 and 6 biological replicates; Mann-Whitney test). (D) Flow cytometric analysis of isolated populations in the spleen for B cell developmental stages T1 (B220^+^AA4.1^+^CD23^-^IgM^+^), T2 (B220^+^AA4.1^+^CD23^+^IgM^+^), and T3 (B220^+^AA4.1^+^CD23^+^IgM^-^). (E) Flow cytometric analysis of isolated populations in the spleen for marginal zone (MZ) B cells (B220^+^CD93^-^CD23^-^IgM^+^) and follicular (FO) B cells (B220^+^CD93^-^CD23^+^IgM^+^). (F) Quantification of splenic populations in (D, E) (n = 7 and 6 biological replicates; Mann-Whitney test). All scatter plots are presented as mean ± SEM.

### Both Orai3 and Orai1 are essential components of SOCE in primary B cells

To determine whether Orai3 regulates Ca^2+^ signals and function of primary B cells, we generated B cell specific Orai3 knockout mice (*Orai3*^*fl/fl*^ *Mb1*^*cre/+*^) and B cell specific double Orai1/Orai3 knockout mice (*Orai1*/*Orai3*^*fl/fl*^ *Mb1*^*cre/+*^). Primary B cells isolated from spleens of Orai1, Orai3, and Orai1/Orai3 knockout mice showed near complete ablation of their respective *Orai* isoform mRNA with no compensatory changes in *Orai2* mRNA expression (**Fig. 5A**). Depletion of ER Ca^2+^ stores with thapsigargin demonstrated that SOCE was significantly reduced (by ∼69%) in Orai1 knockout B cells (as shown in **Fig. 3F, G**), and this remaining Ca^2+^ entry was further reduced in Orai1/Orai3 double knockout B cells (by ∼83%; **Fig. 5B, C**). However, SOCE in single Orai3 knockout B cells was ∼102% of control, which was not significantly different from control B cells.

**Figure 5.**
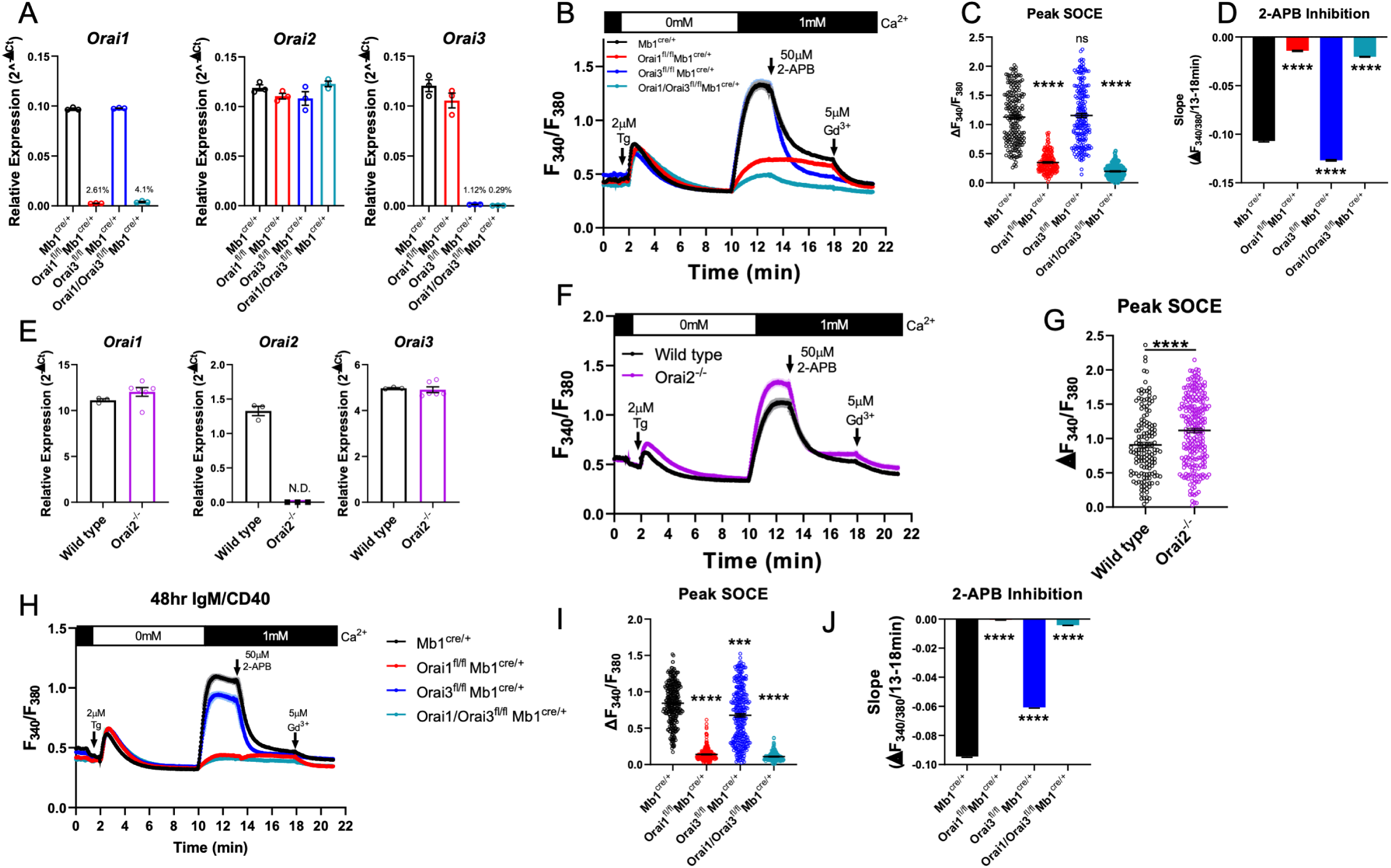
Orai1 and Orai3 are essential components of SOCE in primary B cells. (A) Quantitative RT-PCR of *Orai1, Orai2*, and *Orai3* mRNA in negatively isolated B cells from B cell-specific Orai knockout mice (n = 3 biological replicates per genotype). (B) Measurement of SOCE in naïve B cells with Fura2 upon store depletion with 2µM thapsigargin in 0mM Ca^2+^ followed by re-addition of 1mM Ca^2+^ to the external bath solution. Subsequently, SOCE was inhibited with the addition of 50µM 2-APB at 13 minutes followed by 5µM Gd^3+^ at 18 minutes. (C) Quantification of peak SOCE in (B) (from left to right n = 200, 200, 199, and 149 cells; Kruskal-Wallis test with multiple comparison to Mb1^cre/+^). (D) Quantification of the rate of 2-APB inhibition from 13-18 minutes. (E) Quantitative RT-PCR of *Orai1, Orai2*, and *Orai3* mRNA in negatively isolated B cells from wild-type and Orai2^-/-^ mice (n = 3 and 6 biological replicates). (F) Measurement of SOCE in naïve B cells with Fura2 from wild-type and *Orai2*^*-/-*^ mice. (G) Quantification of peak SOCE in (F) (n = 147 and 240 cells; Mann-Whitney test). (H) Measurement of SOCE in B cells stimulated for 48hr with anti-IgM + anti-CD40. (I) Quantification of peak SOCE in (H) (from left to right n = 287, 288, 290, and 172 cells; Kruskal-Wallis test with multiple comparisons to Mb1^cre/+^). (J) Quantification of the rate of 2-APB inhibition from (H). All scatter plots are presented as mean ± SEM. For all figures, ***p<0.001; ****p<0.0001; ns, not significant.

Orai channel isoforms demonstrate distinct pharmacological profiles and sensitivities to various CRAC channel modifiers (Bird & Putney, 2018; Zhang *et al*, 2020). One of the most extensively utilized CRAC channel modifier is 2-aminoethyldiphenyl borate (2-APB), which at high (25-50µM) concentrations strongly inhibits Orai1, partially inhibits Orai2, and potentiates Orai3 channel activity (DeHaven *et al*, 2008; Zhang *et al*., 2020). After allowing Ca^2+^ entry to plateau, 50µM 2-APB was added, followed by the addition of 5µM gadolinium (Gd^3+^) which potently blocks all Orai isoforms (Yoast *et al*., 2020b; Zhang *et al*., 2020). In wild-type B cells, 2-APB led to a gradual inhibition of SOCE over the course of 5 minutes, which completely returned to baseline following the addition of Gd^3+^. Interestingly, the remaining SOCE in Orai1 knockout B cells showed essentially no inhibition by 2-APB but was strongly inhibited by Gd^3+^. Furthermore, SOCE in Orai3 knockout B cells was inhibited at a significantly faster rate by 2-APB compared to wild-type B cells and was not further inhibited by Gd^3+^ (**Fig. 5B, D**). The small amount of SOCE remaining in Orai1/Orai3 double knockout cells was reduced to baseline following the addition of 2-APB. This remaining SOCE in Orai1/Orai3 double knockout B cells prompted us to measure SOCE in B cells isolated from global Orai2 knockout mice (*Orai2*^*-/-*^). B cells from Orai2^-/-^ mice showed complete loss of *Orai2* mRNA with no apparent compensation in *Orai1* or *Orai3* mRNA (**Fig. 5E**). SOCE stimulated by thapsigargin in B cells from Orai2^-/-^ mice was enhanced by comparison to B cells from wild-type littermate controls (**Fig. 5F, G**), suggesting that, as was shown in T cells(Vaeth *et al*., 2017b), Orai2 is also a negative regulator of SOCE in B cells. Thus, these experiments indicate that the remaining SOCE in B cells from Orai3/Orai1 double knockout mice is most likely mediated by Orai2.

Given the modulation of Orai channel isoforms in response to B cell activation, we performed similar Ca^2+^ imaging recordings on primary B cells that were first activated for 48 hours with anti-IgM + anti-CD40 (**Fig. 5H**). While similar trends were observed as in experiments with naïve primary B cells, several exceptions were notable. SOCE in Orai3 knockout B cells was reduced compared to control cells, like data in A20 Orai knockout cell lines (**Fig. 2A, B**). Differences in inhibition by 50µM 2-APB also became less pronounced between control and Orai3 knockout cells (**Fig. 5H-J**). Collectively, these data demonstrate that Orai3 contributes to SOCE as a component of the native CRAC channel in primary B cells.

### SOCE is an essential regulator of NFAT activation in B cells

Ca^2+^ entry through CRAC channels is critical for the activation of multiple NFAT isoforms(Prakriya & Lewis, 2015; Trebak & Kinet, 2019; Vaeth & Feske, 2018). NFAT nuclear translocation requires Ca^2+^ entry specifically through native Orai1, while native Orai2 and Orai3 were incapable of activating NFAT in cells lacking Orai1 despite mediating a Ca^2+^ signal (Yoast *et al*., 2020b). To determine the role of Orai1 in regulating NFAT1 activation in B cells, primary B cells from control and *Orai1*^*fl/fl*^ *Mb1*^*cre/+*^ mice were stimulated with anti-IgM antibodies and native NFAT1 nuclear translocation was evaluated using ImageStream analysis (**Fig. 6A-C**). In unstimulated B cells, colocalization of endogenous NFAT1 with nuclear DAPI staining was relatively low, and this colocalization increased three-fold following anti-IgM stimulation (**Fig. 6A-C**). However, this anti-IgM mediated NFAT1 translocation was significantly reduced in B cells from *Orai1*^*fl/fl*^ *Mb1*^*cre/+*^ mice (**Fig. 6B, C**).

**Figure 6.**
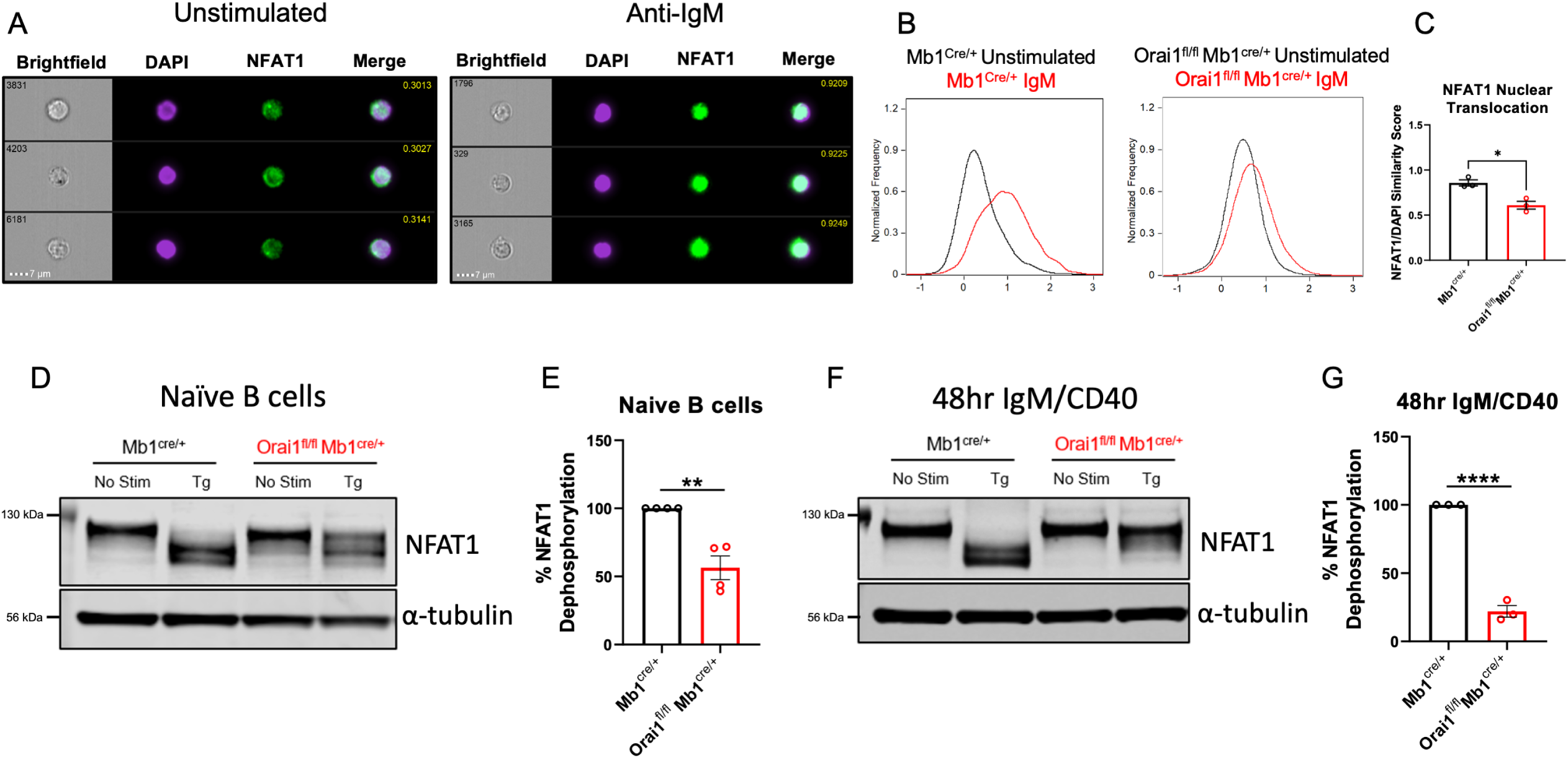
Orai1 is a critical regulator of NFAT activation in naïve and activated B cells. (A) Representative Imagestream images following intracellular staining for NFAT1 and DAPI in naïve B cells from *Mb1*^*cre/+*^ mice before and after 20μg/mL anti-IgM stimulation for 15 minutes. Merge image indicates similarity score co-localization between NFAT1/DAPI. (B) Histograms of NFAT1/DAPI similarity scores before (black trace) and after (red trace) anti-IgM stimulation in naïve B cells from *Mb1*^*cre/+*^ and *Orai1*^*fl/fl*^ *Mb1*^*cre/+*^ mice. (C) Quantification of similarity scores following anti-IgM stimulation in (B) (n = 3 biological replicates for each; unpaired T-test). (D) Western blot analysis of NFAT1 and α-tubulin in naïve B cells isolated from *Mb1*^*cre/+*^ and *Orai1*^*fl/fl*^ *Mb1*^*cre/+*^ mice. B cells were left unstimulated or treated with 2µM thapsigargin (Tg) for 15 minutes before harvesting. (E) Quantification of NFAT1 dephosphorylation in (D) (n = 4 biological replicates for each; Mann-Whitney test). (F) Western blot analysis of NFAT1 and α-tubulin in B cells stimulated for 48 hours with anti-IgM + anti-CD40. (G) Quantification of NFAT1 dephosphorylation in (F) (n = 3 biological replicates for each; Mann-Whitney test). All scatter plots are presented as mean ± SEM. For all figures, *p<0.05; **p<0.01; ****p<0.0001.

We used another complimentary biochemical protocol to assess the nuclear translocation of native NFAT1 in response to stimulation with thapsigargin. After B cell stimulation, proteins were harvested and processed for Western blotting using a specific NFAT1 antibody (**Fig. 6D-G**). In unstimulated samples, NFAT1 appears as a single band corresponding to its highly phosphorylated, non-activated state (**Fig. 6D, F**). Stimulation of naïve B cells with thapsigargin, which completely empties ER stores and maximally activates SOCE, led to a complete shift of the phosphorylated single band into lower molecular weight bands (**Fig. 6D, E**, *left*) and this shift in NFAT1 molecular weight was inhibited by 43.6% in Orai1-deficient B cells (**Fig. 6D, E**, *right*). Given the dynamic expression of each Orai isoform following B cell activation (**Fig. 1G-I**), we also evaluated NFAT1 activation in B cells stimulated for 48-hours with anti-IgM + anti-CD40. While thapsigargin stimulation also led to a complete shift in NFAT1 molecular weight in activated control B cells, this shift was reduced by 78% in activated B cells from *Orai1*^*fl/fl*^ *Mb1*^*cre/+*^ mice (**Fig. 6F, G**). Similar results were obtained when we analyzed NFAT2 isoform dephosphorylation in naïve B cells (**Fig. 6-Fig. Supplement 1A**). Furthermore, when naïve B cells were stimulated with anti-IgM (instead of thapsigargin), we observed similar trends of NFAT1 and NFAT2 dephosphorylation, although this dephosphorylation was not as robust as with thapsigargin (**Fig. 6-Fig. Supplement 1A**). Nevertheless, this anti-IgM mediated dephosphorylation of NFAT1 and NFAT2 was inhibited in B cells from *Orai1*^*fl/fl*^ *Mb1*^*cre/+*^ mice (**Fig. 6-Fig. Supplement 1A**).

We also evaluated NFAT1 activation in response to thapsigargin in both naïve B cells (**Fig. 6-Fig. Supplement 1B**) and activated B cells (48 hours with anti-IgM + anti-CD40; **Fig. 6-Fig. Supplement 1C**) isolated from either *Orai3*^*fl/fl*^ *Mb1*^*cre/+*^ or *Orai3/Orai1*^*fl/fl*^ *Mb1*^*cre/+*^ mice. Loss of Orai3 alone from naïve B cells had no effect on NFAT1 activation, while naïve B cells isolated from double Orai1/Orai3 knockout cells showed reductions in NFAT1 activation comparable to B cell isolated from single Orai1 knockout mice (**Fig. 6-Fig. Supplement 1B**). Importantly, this impairment of NFAT1 activation was further exacerbated in activated B cells isolated from Orai1 knockout and Orai1/Orai3 double knockout mice (**Fig. 6-Fig. Supplement 1C**). Collectively, these data reveal that Orai1 plays a more prominent role in the activation of multiple NFAT isoforms in activated B cells compared to naïve cells.

### Orai3 and Orai1 coordinate for efficient B cell proliferation

BCR-induced Ca^2+^ signals that are sustained through SOCE are critical for the activation of gene programs that regulate proliferation and apoptosis (Berry *et al*., 2020; Matsumoto *et al*., 2011). Interestingly, suboptimal proliferation and survival of B cells in response to BCR stimulation can largely be rescued through the addition of secondary co-stimulatory signals (e.g. CD40 or TLR stimulation) (Berry *et al*., 2020; Matsumoto *et al*., 2011; Tang *et al*, 2017). To understand how Ca^2+^ signals downstream of Orai isoforms regulate B cell expansion, primary B cells were labeled with cabroxyfluorescein diacetate succinimidyl ester (CFSE) and cell divisions tracked in response to multiple stimulation conditions (**Fig. 7A**). BCR stimulation with anti-IgM alone resulted in multiple rounds of cell division in most wild-type cells (87.7%) with 28.6% of viable cells at 72 hours post stimulation (**Fig. 7B, C**, *black*). B cell viability at 72 hours was dramatically increased to 67.4% when cells were co-stimulated with anti-IgM + anti-CD40, with an increase in the number of cells undergoing several cycles of cell division. Surprisingly, following anti-IgM stimulation Orai1-deficient B cells showed a similar percentage of proliferating cells (83.1%) as controls, with a moderate reduction in cell viability to 20.9% (**Fig. 7B, C**, *red*). Interestingly, while anti-IgM mediated proliferation of Orai3-deficient B cells was comparable to controls (90.3%), their viability was 34.8% (**Fig. 7B, C**, *blue*). Importantly, combined deletion of Orai1 and Orai3 resulted in a substantial reduction in the percentage of both viable (10%) and proliferating (60.8%) cells in response to anti-IgM stimulation (**Fig. 7B, C**, *teal*). These defects in survival and proliferation of Orai1/Orai3-deficient B cells were partially rescued when cells were co-stimulated with anti-IgM + anti-CD40, albeit to a lesser extent than control and single Orai knockout B cells. Similar percentages of viability and proliferation were observed in all experimental groups when B cells were stimulated with either anti-CD40 or LPS. We also analyzed the proliferation and survival of B cells isolated from Orai2^-/-^ mice and their wildtype littermates exposed to the same stimuli. We found only marginal or no effects of Orai2 knockout on B cell proliferation and survival (**Fig. 7-Fig. Supplement 1A, B**). Furthermore, we performed RNA sequencing on B cells isolated from control *Mb1*^*cre/+*^ and *Orai3*/*Orai1*^*fl/fl*^ *Mb1*^*cre/+*^ mice after stimulation with anti-IgM for 24 hours. In agreement with our proliferation and survival data, gene set enrichment analysis (GSEA) showed significant downregulation of pathways that govern survival and cell cycle progression in Orai1/Orai3-deficient B cells (**Fig. 7D, E**). Together, these findings reveal that the coordinated functions of both Orai1 and Orai3 are critical for B cell survival and proliferation.

**Figure 7.**
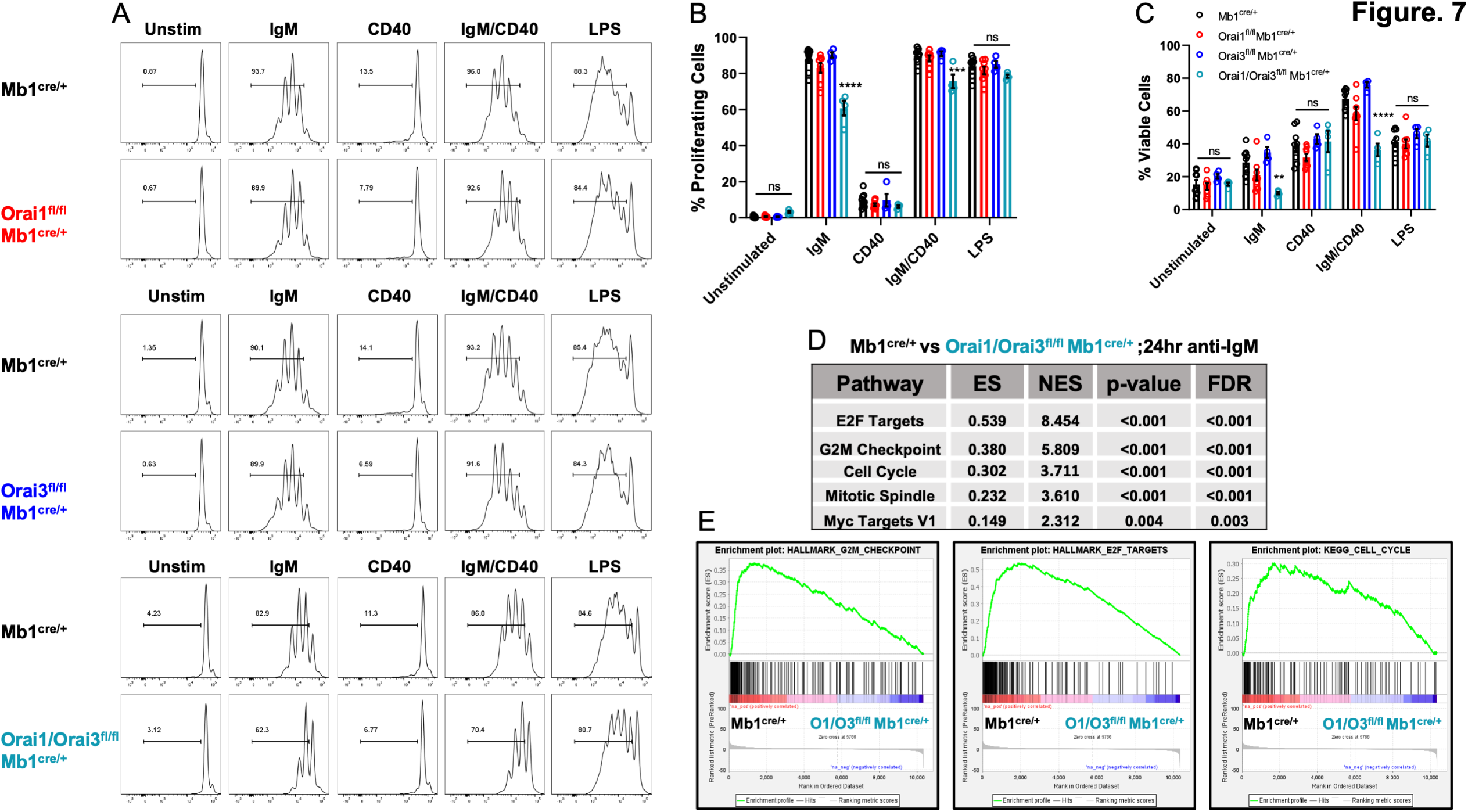
Orai1 and Orai3 regulate B cell proliferation and survival. (A) Measurement of B cell proliferation by tracking CFSE dilution. B lymphocytes from control, Orai1, Orai3, and Orai1/Orai3 knockout mice were loaded with CFSE (3μM) and stimulated with anti-IgM (20μg/mL), anti-CD40 (10μg/mL), anti-IgM + anti-CD40, or LPS (10μg/mL). CFSE dilution was determined 72 hours after stimulation for all conditions. (B) Quantification of the percentage of proliferating cells for each condition in (A) (from left to right n = 9, 8, 4, and 4 biological replicates; one-way ANOVA with multiple comparisons to Mb1^cre/+^). CFSE dilution gate is drawn relative to unstimulated controls. (C) Quantification of the percentage of viable cells for each condition in (A) as determined by a Live/Dead viability dye (from left to right n = 9, 8, 4, and 4 biological replicates; one-way ANOVA with multiple comparisons to Mb1^cre/+^). (D) Top KEGG pathways showing differential expression from RNA-sequencing analysis of B cells from *Mb1*^*cre/+*^ vs *Orai1/Orai31*^*fl/fl*^ *Mb1*^*cre/+*^ mice stimulated for 24 hours with anti-IgM (20μg/mL) (n = 3 biological replicates for each). (E) Gene set enrichment analysis (GSEA) plots of the G2M Checkpoint, E2F Targets, and Cell Cycle gene sets. All scatter plots are presented as mean ± SEM. For all figures, **p<0.01; ***p<0.001; ****p<0.0001; ns, not significant.

### Orai3 and Orai1 are essential regulators of B cell metabolism

Metabolic reprogramming results in dramatically enhanced OXPHOS and remodeling of the mitochondrial network (Akkaya *et al*., 2018; Waters *et al*, 2018). Indeed, gene set enrichment analysis (GSEA) of B cells stimulated for 24 hours with anti-IgM identified the Oxidative Phosphorylation gene set as one of the most highly enriched pathways relative to unstimulated controls (**Fig. 8A**). Additionally, anti-IgM stimulated B cells demonstrate a significant increase in the total number of mitochondria per cell compared to naïve B cells as determined by transmission electron microscopy (TEM; **Fig. 8B, C**). However, this increase in mitochondrial mass triggered by B cell activation was not affected by depletion of Orai1 from B cells, as determined by Mitotracker Green staining (**Fig. 8D**). We measured OCR and ECAR in B cells isolated from control and B cell specific Orai knockout mice after B cell stimulation with anti-IgM for 24 hours (**Fig. 8E**). Anti-IgM stimulation of wild-type B cells led to a robust increase in both OCR and ECAR (**Fig. 8F**). Furthermore, Anti-IgM stimulation led to an increase in both basal and maximal respiration of wild-type B cells (**Fig. 8E-H**). Interestingly, the increase in OCR and ECAR following BCR activation was significantly blunted in Orai1-deficient B cells, and further reduced in Orai1/Orai3-deficient B cells (**Fig. 8E-H**). Loss of Orai3 alone only partially reduced glycolytic flux and basal respiration but did not affect maximal respiration (**Fig. 8E-H**). Previous research has shown that loss of SOCE in the chicken DT40 B cell line impaired mitochondrial metabolism by reducing CREB-mediated expression of the mitochondrial calcium uniporter (MCU) (Shanmughapriya *et al*, 2015). We observed no differences in CREB phosphorylation on Ser133 (a surrogate for CREB activation) or MCU expression in primary Orai1-deficient B cells from *Orai1*^*fl/fl*^ *Mb1*^*cre/+*^ mice (**Fig. 8-Fig. Supplement 1A**). Likewise, we observed no differences in MCU expression in mouse A20 B cells lacking either Orai1, Orai3, or both Orai1 and Orai3 (**Fig. 8-Fig. Supplement 1B**). Both naïve and activated (24 hours with anti-IgM) B cells from *Orai1*^*fl/fl*^ *Mb1*^*cre/+*^ mice showed no obvious changes in expression of different components of the electron transport chain (**Fig. 8-Fig. Supplement 1C, D**).

**Figure 8.**
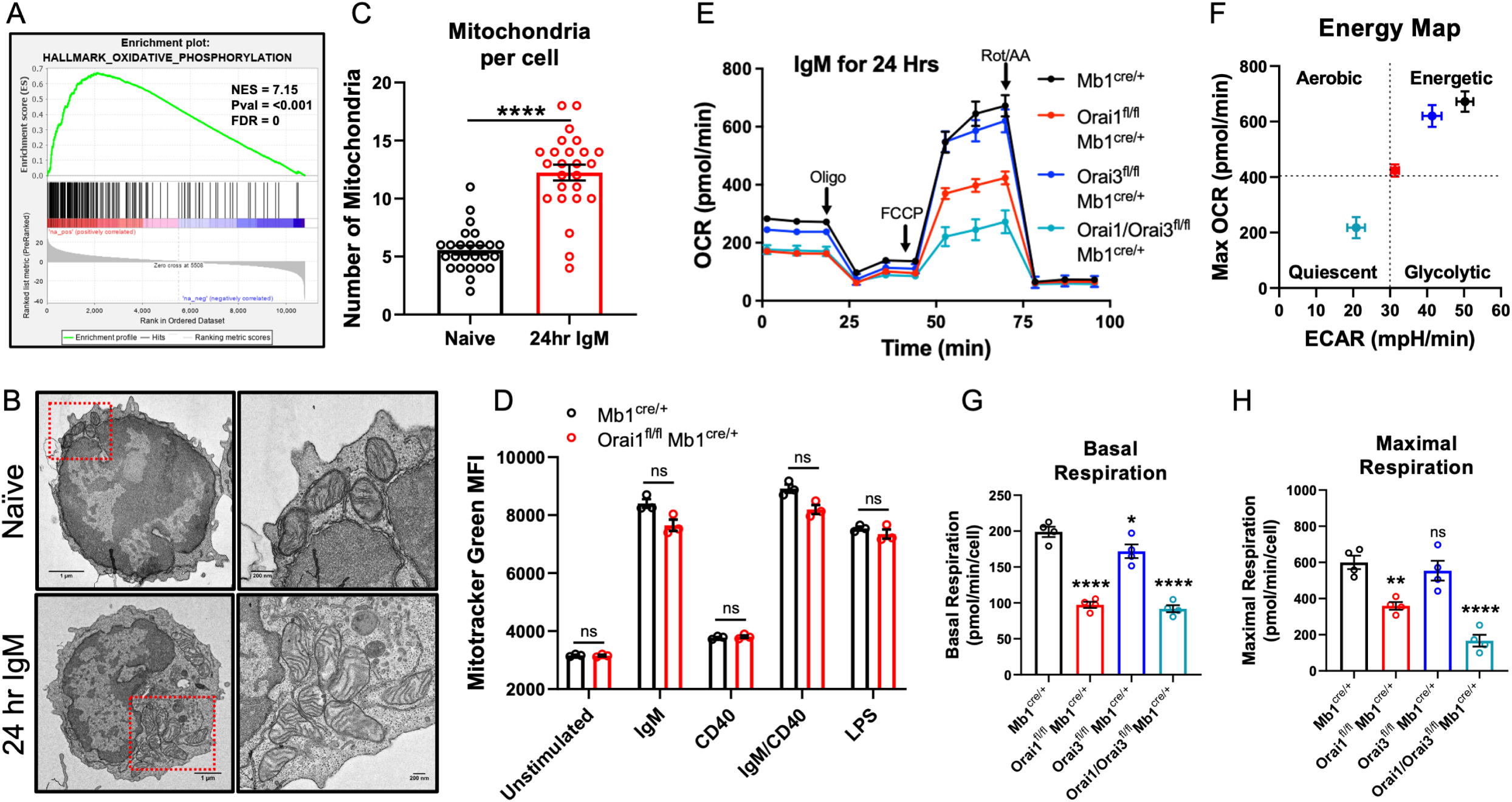
Orai1 and Orai3 are critical for B cell mitochondrial respiration. (A) Gene set enrichment analysis (GSEA) of the KEGG Oxidative Phosphorylation gene set in B cells stimulated for 24 hours with anti-IgM (20μg/mL) relative to unstimulated controls. (B) Representative TEM images of B cells from *Mb1*^*cre/+*^ mice. Shown are naïve, unstimulated B cells (top) and B cells stimulated for 24 hours with anti-IgM (bottom). (C) Quantification of total mitochondria per cell in unstimulated B cells and B cells stimulated for 24 hours with anti-IgM (n = 25 for each; Mann-Whitney test). (D) Measurement of total mitochondrial content with MitoTracker Green in B cells from *Mb1*^*cre/+*^ and *Orai1*^*fl/fl*^ *Mb1*^*cre/+*^ mice following 24-hour stimulation (n = 3 biological replicates for each; Mann-Whitney test). (E) Measurement of oxygen consumption rate (OCR) in primary B lymphocytes following 24-hour stimulation with anti-IgM (20μg/mL) using the Seahorse Mito Stress Test. (F) Energy map of maximal OCR and ECAR following FCCP addition. (G, H) Quantification of basal (G) and maximal (H) respiration from Seahorse traces in (E) (n = 4 biological replicates for each genotype; one-way ANOVA with multiple comparisons to Mb1^cre/+^). All scatter plots and Seahorse traces are presented as mean ± SEM. For all figures, **p<0.01; ****p<0.0001; ns, not significant.

To further investigate how Orai-mediated Ca^2+^ signaling regulates B cell metabolism, we profiled polar metabolites utilizing liquid chromatography followed by mass spectrometry in control and Orai1/Orai3 deficient B cells (**Fig. 9**). Isolated B cells from each genotype were either unstimulated or stimulated for 24 hours with anti-IgM alone, anti-IgM + anti-CD40, anti-IgM + the calcineurin inhibitor FK506, or anti-IgM + the CRAC channel inhibitor GSK-7975A. We utilized GSK-7975A as we recently demonstrated that this compound inhibited all Orai isoforms compared to differential effects on Orai isoforms with other common SOCE inhibitors like Synta66 (Zhang *et al*., 2020). Stimulation of control B cells with anti-IgM led to a significant increase of glycolytic and TCA cycle metabolites along with most non-essential amino acids. This effect was further enhanced with anti-IgM + anti-CD40 co-stimulation (**Fig. 9A; Fig. 9-Fig. Supplement 1A**). Inclusion of FK506 or GSK-7975A strongly blunted the effects of anti-IgM activation as overall metabolic profiles remained like unstimulated cells (**Fig. 9A**). Importantly, this upregulation in polar metabolites upon B cell activation was significantly reduced in Orai1/Orai3-deficient B cells with either anti-IgM stimulation or anti-IgM + anti-CD40 co-stimulation (**Fig. 9B, C; Fig. 9-Fig. Supplement 1B**). Of note, upregulation of most polar metabolites from Orai1/Orai3 knockout B cells co-stimulated with anti-IgM + anti-CD40 typically reached levels comparable to those of control B cells stimulated with anti-IgM alone. The inhibitory effects of FK506 and GSK-7975A on the metabolite status of wildtype B cells were comparable to those of Orai1/Orai3-deficient B cells (**Fig. 9A, Fig. 9-Fig. Supplement 1A**). Collectively, these results reveal that SOCE mediated through Orai1/Orai3 channels is crucial to the metabolic reprogramming of B lymphocytes.

**Figure 9.**
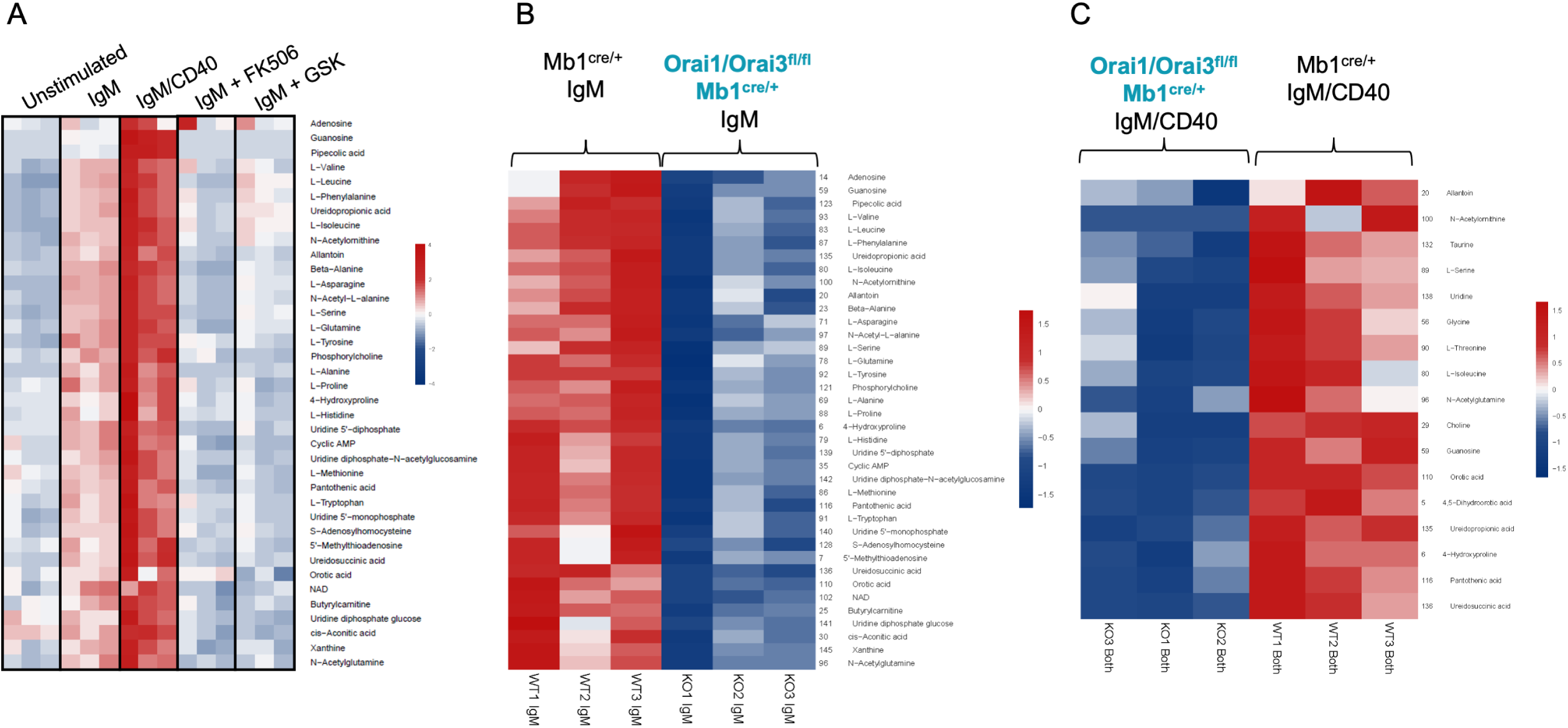
Orai1/Orai3-mediated SOCE-calcineurin-NFAT pathway regulates B cell metabolism. (A) Analysis of polar metabolites in B cells from *Mb1*^*cre/+*^ mice utilizing liquid chromatography followed by mass spectrometry. B cells were either unstimulated or stimulated for 24-hours with anti-IgM, anti-IgM + anti-CD40, anti-IgM with 1µM FK506, or anti-IgM with 10µM GSK-7975A. (B, C). Heat maps of statistically significant polar metabolites in B cells from *Mb1*^*cre/+*^ and *Orai1/Orai31*^*fl/fl*^ *Mb1*^*cre/+*^ mice following (B) 24-hour anti-IgM stimulation or (C) 24-hour anti-IgM + anti-CD40 stimulation. (n = 3 biological replicates for each condition).

## Discussion

Given the essential role of B lymphocytes in driving humoral immunity against foreign pathogens, a comprehensive understanding of the molecular pathways that govern their development, differentiation, and effector functions is critical for future targeted therapies. One of the earliest signaling events upon crosslinking of the BCR is a biphasic increase in intracellular Ca^2+^ concentrations (Baba & Kurosaki, 2016). Early landmark studies established that multiple Ca^2+^ dependent transcription factors display unique activation requirements by relying on either ER Ca^2+^ release (e.g. JNK, NF-κB) or sustained Ca^2+^ signals driven by SOCE (NFAT) (Dolmetsch *et al*, 1997; Healy *et al*, 1998). While the SOCE-calcineurin-NFAT pathway is well established in the context of B cell effector function, recent reports have shed light on the mechanisms by which SOCE also regulates NF-κB activation and its downstream target genes (Berry *et al*., 2020; Berry *et al*., 2018). These two BCR-activated signaling pathways are sustained by Ca^2+^ entry through CRAC channels and synergize with one another to activate a series of Ca^2+^-regulated checkpoints that determine B cell survival, entry into the cell cycle, and proliferation (Akkaya *et al*., 2018; Berry *et al*., 2020). Unlike recent findings that have established Orai1 and Orai2 as the major Orai isoforms mediating SOCE in T cells (Vaeth *et al*., 2017b), the composition of the native CRAC channel in B cells remained, until now, unclear.

Our results herein establish that B cell activation through BCR stimulation alone or co-stimulation with secondary signals like CD40 or TLR ligands significantly enhanced metabolic activity. Concurrently, these stimulation conditions drive dynamic changes in the expression of each Orai isoform. Interestingly, we observe that robust B cell activation with BCR and CD40 co-stimulation results in upregulation of both Orai1 and Orai3, and downregulation of Orai2. While Orai1 has previously been shown to contribute to the majority of SOCE in B cells under conditions of maximal ER Ca^2+^ depletion (Gwack *et al*., 2008; McCarl *et al*., 2010), we show using B-cell specific Orai1 knockout mice that Orai1 is dispensable for maintaining cytosolic Ca^2+^ oscillations in response to BCR crosslinking. In agreement with our previous results in HEK293 cells (Yoast *et al*., 2020b), we also show that Orai1 is an essential regulator of NFAT1 and NFTA2 isoforms in B cells and that its role becomes more prominent for NFAT induction in activated B cells compared to naïve unstimulated B cells. By generating CRISPR/Cas9 B cell lines and B-cell specific knockout mice lacking Orai1 or Orai3 individually and in combination, we found that Orai3 is an essential component of the native SOCE pathway in B cells. Importantly, combined loss of Orai1 and Orai3, but not either isoform alone, led to a significant reduction in both B cell proliferation and survival. Similarly, we reveal that SOCE through Orai1/Orai3 is critical for the metabolic reprogramming of B cells by regulating mitochondrial metabolism and the flux of polar metabolites in response to B cell activation. This shift in metabolic profiles in response to B cell activation could be neutralized by inhibition of either calcineurin or CRAC channels with FK506 or GSK-7975A respectively, suggesting that this metabolic flux is driven through SOCE and NFAT dependent mechanisms. Collectively, these data demonstrate that both Orai1 and Orai3 are critical in mediating SOCE and shaping cytosolic Ca^2+^ signaling in B cells.

While our data has provided evidence for the contribution of Orai3 to native CRAC channels in B cells, its contribution in other immune cell populations is less clear. Earlier studies investigating human effector T cells demonstrated that Orai3 becomes upregulated in response to oxidative stress, which may act as a potential mechanism to maintain a threshold of SOCE and T cell function in various inflammatory conditions (Bogeski *et al*, 2010). In support of this model, new findings have demonstrated that Orai3 expression is increased in CD4^+^ T cells from patients with rheumatoid arthritis and psoriatic arthritis and that silencing of Orai3 reduces tissue inflammation in a human synovium adoptive transfer model (Ye *et al*, 2021). Interestingly, these effects were proposed to be through the store-independent function of Orai3 by mediating Ca^2+^ influx in response to arachidonic acid or its metabolite, leukotrieneC_4_ (Thompson *et al*, 2013; Zhang *et al*, 2015; Zhang *et al*, 2013; Zhang *et al*, 2018; Zhang *et al*, 2014). In contrast, combined loss of Orai1 and Orai2 in murine T cells led to a near complete ablation of SOCE and substantially impaired T cell function (Vaeth *et al*., 2017b). Whether Orai3 contributes to the small, residual amount of SOCE in Orai1/Orai2-deficient T cells and/or mediates store-independent signaling functions in murine T and B cells is unknown. Furthermore, differences in Orai channel contributions are apparent between mice and humans, as SOCE and CRAC currents are completely inhibited in T cells from patients with LoF mutations in *Orai1*, while SOCE is only partially inhibited in Orai1-deficient murine T cells (Feske *et al*., 2006; Lian *et al*, 2018; McCarl *et al*., 2009; Vig *et al*, 2008). Curiously, B cells from *Orai1* LoF mutation patients still retain residual SOCE, suggesting that another Orai isoform may play a more prominent role in human B cells than in T cells (Feske *et al*, 2001), in agreement with our data herein in mouse B cells. Thus, the composition of native CRAC channels among lymphocytes appears cell-type and context specific.

While SOCE is substantially reduced in Orai1/Orai3-deficient B cells, defects in their ability to proliferate and survive in response to antigenic stimulation are not as severe as the phenotype observed in STIM1/STIM2-deficient B cells (Berry *et al*., 2020; Matsumoto *et al*., 2011). Our data clearly establish cooperativity between Orai1 and Orai3 channels in controlling B cell survival and proliferation, but we cannot exclude a potential role for Orai2 in the regulation of SOCE in B cells. Quite the opposite, we show that Orai2 is functional in B cells by documenting that B cells from Orai2^-/-^ mice have enhanced SOCE. This suggests that Orai2 acts as a negative regulator of SOCE in B cells as was shown for T cells (Vaeth *et al*., 2017b), and that the remaining SOCE activity in B cells from Orai1/Orai3 double knockout mice is likely mediated by Orai2. Nevertheless, development of Orai triple knockout mice and additional Orai knockout pairs is needed to further clarify this issue. Similarly, how SOCE and the composition of the native CRAC channel varies among different B cell subsets and during pathological conditions is completely unknown. Indeed, earlier studies have demonstrated that effector populations like germinal center B cells display unique metabolic and Ca^2+^ signaling requirements (Khalil *et al*, 2012; Luo *et al*, 2018). Our data also suggest that expression of each Orai isoform is dynamic in response to B cell activation, similar to the case of naïve vs effector T cells (Vaeth *et al*., 2017b). Taken together, our results uncover a novel role of Orai3 in the immune system and further clarify the complex signaling functions of SOCE in B lymphocytes.

## Methods

### Mice

All animal experiments were carried out in compliance with the Institutional Animal Care and use Committee (IACUC) guidelines of the Pennsylvania State University College of Medicine and in accordance with the ARRIVE guidelines for reporting animal research. All mice were housed under specific pathogen-free conditions and experiments were performed in accordance with protocols approved by the IACUC of the Pennsylvania State University College of Medicine. Mb1-Cre mice(Hobeika *et al*, 2006) were obtained from The Jackson Laboratory. *Orai1*^*fl/fl*^ mice(Ahuja *et al*, 2017) were obtained from Dr. Paul Worley (Johns Hopkins University). *Orai3*^*fl/fl*^ mice were generated by our laboratory through the MMRC at University of California Davis. A trapping cassette was generated including “SA-βgeo-pA” (splice acceptor-beta-geo-polyA) flanked by Flp-recombinase target “FRT” sites, followed by a critical *Orai3* coding exon flanked by Cre-recombinase target “loxP” sites. This cassette was inserted within an intron upstream of the *Orai3* critical exon, where it tags the *Orai3* gene with the lacZ reporter. This creates a constitutive null *Orai3* mutation in the target *Orai3* gene through efficient splicing to the reporter cassette resulting in the truncation of the endogenous transcript. Mice carrying this allele were bred with FLP deleter C57BL/6N mice to generate the *Orai3*^*fl/fl*^ mouse. All experiments were performed with 8–12-week-old age and sex matched mice.

### Cell Culture

Parental A20 cells were purchased from ATCC (Catalog # TIB-208) and cultured in RPMI 1640 with L-glutamine supplemented with 10% fetal bovine serum and 1X Antibiotic-Antimycotic. Naïve splenic B lymphocytes were purified by negative selection using the EasySep Mouse B Cell Isolation Kit (STEMCELL Technologies). Primary B cells from each transgenic mouse line were cultured in RPMI 1640 with L-glutamine supplemented with 10% fetal bovine serum, 1X GlutaMAX, 1X Penicillin-Streptomycin Solution, sodium pyruvate (1mM), 2-ME (5uM), and HEPES (10mM). Primary lymphocytes were stimulated with F(ab’)_2_ Fragment Goat Anti-Mouse IgM antibody (Jackson ImmunoResearch, 20µg/mL), anti-mouse CD40 antibody (BioXCell, 10µg/mL), or LPS (Sigma-Aldrich, 10µg/mL). All cell lines and primary lymphocytes were cultured at 37°C in a humidified incubator with 5% CO_2_.

### Generation of A20 Orai CRISPR/Cas9 knockout cells

For generation of A20 Orai knockout clones, we used a similar strategy to our previous studies(Emrich *et al*., 2021; Yoast *et al*, 2021; Yoast *et al*., 2020b; Zhang *et al*, 2019). Briefly, two gRNAs targeting the beginning and end of the mouse *Orai1* or *Orai3* coding region were cloned into two fluorescent vectors (pSpCas9(BB)-2A-GFP and pU6-(BbsI)_CBh-Cas9-T2AmCherry; Addgene). For *mOrai1* knockout, the gRNA sequences are the following: mOrai1n: 5’-GCCTTCGGATCCGGTGCGTC-3’; mOrai1c: 5’-CACAGGCCGTCCTCCGGACT-3’. For *mOrai3* knockout, the gRNA sequences are the following: mOrai3n: 5’- GCGTCCGTAACTGTTCCCGC-3’; mOrai3c: 5- GAAGGAGGTCTGTCGATCCC-3’. A20 cells were electroporated with both N- and C-terminal gRNA combinations with an Amaxa Nucleofector II and single cells with high GFP and mCherry expression were sorted at one cell per well into 96-well plates 24 hours after transfection using a FACS Aria SORP Cell Sorter. Individual clones were obtained and genomic DNA tested using primers targeting the N- and C-terminal gRNA cut sites to resolve wild-type and knockout PCR products. Knockout PCR products were cloned into pSC-B-amp/kan with the StrataClone Blunt PCR Cloning Kit (Agilent) and Sanger sequenced to determine the exact deletion. Knockout clones were also confirmed for absence of mRNA with qRT-PCR and functionally through Ca^2+^ imaging experiments.

### Fluorescence Imaging

A20 cell lines and primary B cells were seeded onto poly-L-lysine (Sigma-Aldrich) coated coverslips. Coverslips were mounted in Attofluor cell chambers (Thermo Scientific) and loaded with 2µM Fura-2AM (Molecular Probes) in a HEPES-buffered saline solution (HBSS) containing 120mM NaCl, 5.4mM KCl, 0.8mM MgCl_2_, 1mM CaCl_2_, 20mM HEPES, 10mM D-glucose, at pH 7.4 for 30 minutes at room temperature. Following Fura-2 loading, cells were washed 3 times with HBSS and mounted on a Leica DMi8 fluorescence microscope. Fura-2 fluorescence was measured every 2 seconds by excitation at 340nm and 380nm using a fast shutter wheel and the emission at 510nm was collected through a 40X fluorescence objective. Fluorescence data was collected from individual cells on a pixel-by-pixel basis and processed using Leica Application Suite X. All cytosolic Ca^2+^ concentrations are presented as the ratio of F340/F380.

### Flow cytometry

Spleens were processed into single-cell suspensions and stained using the following antibodies: B220-BV605 (RA3-6B2, Biolegend), CD86-PE/Cy5 (GL-1, Biolegend), MHC-II-PE/Cy7 (M5/114.15.2, Biolegend), Orai1 rabbit polyclonal (antibody generated by Stefan Feske, NYU), CD3e-PE (145-2C11, BD Biosciences), CD8a-V500 (53-6.7, BD Biosciences), CD4-BB700 (RM4-5, BD Biosciences), APC–anti-CD24 (HSA; M1/69), FITC–anti-CD23 (B3B4), PE–anti-IgM (eB121-15F9, eBioscience), APC–anti-CD93 (AA4.1, eBioscience), PE–Cy5-streptavidin (Biolegend), and Pacific blue–anti-B220 (RA3-6B2, Biolegend). All staining was performed in FACS buffer (DPBS, 2% FBS, 1mM EDTA) for 30 minutes at 4°C. Prior to surface staining, all cells were stained with eBioscience Fixable Viability Dye eFluor 780 (Thermo Fisher) and anti-CD16/32 Fc block (2.4G2, Tonbo Biosciences). Measurement of mitochondrial content was performed using MitoTracker Green (Molecular Probes) and mitochondrial membrane potential performed using TMRE (Molecular Probes). For all CFSE dilution experiments, primary B lymphocytes were stained with 3µM CFSE (C1157, Thermo Fischer) in PBS with 5% FBS for 5 minutes at room temperature, followed by the addition of FBS and two washes in PBS before resuspension in complete RPMI media. CFSE dilution of labeled cells was assessed 72 hours after plating and stimulation. All flow cytometry data was collected on a BD LSR II flow cytometer using FACSDiva software (BD Biosciences) and analyzed with FlowJo 9.9.6 software (Tree Star).

### Analysis of NFAT nuclear translocation

Primary B cells were purified using the EasySep Mouse B Cell Isolation Kit (STEMCELL Technologies) and stimulated with anti-IgM (20µg/mL) for 15 minutes at room temperature in complete RPMI media. Cells were fixed and permeabilized using the Foxp3 / Transcription Factor Staining Buffer Set (eBioscience) following the manufacturer’s instructions. Cells were stained in 1X permeabilization buffer for 1 hour at room temperature with NFAT1-Alexa Fluor 488 (D43B1, Cell Signaling Technology). Nuclei were stained with DAPI prior to acquisition on an Amnis ImageStream X Mark II Imaging Flow Cytometer and analyzed with IDEAS software using the ‘‘Nuclear Localization’’ feature (EMD Millipore).

### Western blot analysis

A20 and primary B lymphocytes were harvested from culture and lysed for 10 minutes in RIPA buffer (150mM NaCl, 1.0% IGEPAL CA-630, 0.5% sodium deoxycholate, 0.1% SDS, 50 mM Tris, pH 8.0; Sigma) containing 1X Halt protease/phosphatase inhibitors (Thermo Scientific). Following lysis, samples were clarified by centrifugation at 15,000xg for 10 minutes at 4°C. Supernatants were collected and protein concentration was determined using the Pierce Rapid Gold BCA Protein Assay Kit (Thermo Scientific). Equal concentrations of protein extract were loaded into 4%–12% NuPAGE BisTris gels (Life Technologies) and transferred to PVDF membranes utilizing the Transblot Turbo Transfer System (Bio-Rad). Membranes were blocked for 1 hour at room temperature in Odyssey Blocking Buffer in TBS (LI-COR) and incubated overnight at 4°C with primary antibody. The following antibodies and dilutions were used: MCU (1:2000; 14997S, Cell Signaling Technology), GAPDH (1:5000; MAB374, Sigma), Total OXPHOS Rodent Antibody Cocktail (1:1000, ab110413, Abcam), NFAT1 (1:1000, 4389S, Cell Signaling Technology), NFAT2 (1:1000, 8032S, Cell Signaling Technology), α-Tubulin (1:5000, 3873S, Cell Signaling Technology), phospho-CREB (Ser133, 1:1000, 9198S, Cell Signaling Technology), and CREB (1:1000, 9104S, Cell Signaling Technology). Membranes were washed with TBST and incubated for 1 hour at room temperature with the following secondary antibodies: IRDye 680RD goat anti-mouse (1:10,000 LI-COR) or IRDye 800RD donkey anti-rabbit (1:10,000 LI-COR). Membranes were imaged on an Odyssey CLx Imaging System (LI-COR) and analysis performed in Image Studio Lite version 5.2 (LI-COR) and ImageJ.

### Quantitative RT-PCR

Total mRNA was isolated from A20 cells and primary B lymphocytes using a RNeasy Mini Kit (Qiagen) following the manufacturer’s instructions. RNA concentrations were measured using a NanoDrop 2000 (Thermo Scientific) and 1µg of DNAse I treated RNA was used with the High-Capacity cDNA Reverse Transcription Kit (Applied Biosystems). cDNA was amplified on a QuantStudio 3 Real-Time PCR System (Applied Biosystems) using PowerUp SYBR Green Master Mix (Applied Biosystems). PCR amplification was performed by initial activation for 2 minutes at 50 °C, followed by a 95 °C 2-minute melt step. The initial melt steps were then followed by 40 cycles of 95 °C for 15 seconds, 60°C for 15 seconds, and 72 °C for 30 seconds. Data were analyzed with the instrument software v1.3.1 (Applied Biosystems) and analysis of each target was carried out using the comparative Ct method.

### Seahorse Extracellular Flux Analysis

Oxygen consumption rates (OCR) and extracellular acidification rates (ECAR) were measured using an XFe24 Extracellular Flux Analyzer (Seahorse Bioscience). A20 cells (0.8×10^6^ per well) and primary B lymphocytes (2×10^6^ per well) were resuspended in XF DMEM pH 7.4 media supplemented with 1mM pyruvate, 2mM glutamine, and 10mM glucose (Mito Stress Test) and plated in poly-L-lysine coated microchamber wells. For the Mito Stress Test, 1.5 μM oligomycin, 2 μM FCCP, and 0.5 μM antimycin/rotenone were utilized. Data were analyzed using the Agilent Seahorse Wave Software and normalized to total protein context per well using the Pierce Rapid Gold BCA Protein Assay Kit (Thermo Scientific).

### Transmission Electron Microscopy

Primary B cells (unstimulated or 24-hour IgM stimulated) were seeded onto poly-L-lysine coated cell culture dishes and fixed with 1% glutaraldehyde in 0.1 M sodium phosphate buffer, pH 7.3. After fixation, the cells were washed with 100 mM Tris (pH 7.2) and 160 mM sucrose for 30 minutes. The cells were washed twice with phosphate buffer (150 mM NaCl, 5 mM KCl, 10 mM Na3PO4, pH 7.3) for 30 minutes, followed by treatment with 1% OsO4 in 140 mM Na3PO4 (pH 7.3) for 1 hour. The cells were washed twice with water and stained with saturated uranyl acetate for 1 hour, dehydrated in ethanol, and embedded in Epon (Electron Microscopy Sciences, Hatfield, PA). Roughly 60 nm sections were cut and stained with uranyl acetate and lead nitrate. The stained grids were analyzed using a Philips CM-12 electron microscope (FEI; Eindhoven, The Netherlands) and photographed with a Gatan Erlangshen ES1000W digital camera (Model 785, 4 k 3 2.7 k; Gatan, Pleasanton, CA).

### RNA Sequencing and Differential Expression Analysis

Total mRNA was isolated from primary B lymphocytes using a RNeasy Mini Kit (Qiagen) following the manufacturer’s instructions. RNA concentrations were quantitated using a NanoDrop 2000 (Thermo Scientific) and library preparation was performed by Novogene. A total amount of 1 µg RNA per sample was used as input material for the RNA sample preparations. Sequencing libraries were generated using NEBNext® Ultra TM RNA Library Prep Kit for Illumina® (NEB, USA) following manufacturer’s recommendations and index codes were added to attribute sequences to each sample. Briefly, mRNA was purified from total RNA using poly-T oligo-attached magnetic beads. Fragmentation was carried out using divalent cations under elevated temperature in NEBNext First Strand Synthesis Reaction Buffer (5X). First strand cDNA was synthesized using random hexamer primer and M-MuLV Reverse Transcriptase (RNase H-). Second strand cDNA synthesis was subsequently performed using DNA Polymerase I and RNase H. Remaining overhangs were converted into blunt ends via exonuclease/polymerase activities. After adenylation of 3’ ends of DNA fragments, NEBNext Adaptor with hairpin loop structure were ligated to prepare for hybridization. To select cDNA fragments of preferentially 150∼200 bp in length, the library fragments were purified with AMPure XP system (Beckman Coulter, Beverly, USA). Then 3 µl USER Enzyme (NEB, USA) was used with size-selected, adaptor-ligated cDNA at 37 °C for 15 min followed by 5 min at 95 °C before PCR. Then PCR was performed with Phusion High-Fidelity DNA polymerase, Universal PCR primers and Index (X) Primer. Lastly, PCR products were purified (AMPure XP system) and library quality was assessed on the Agilent Bioanalyzer 2100 system. The clustering of the index-coded samples was performed on a cBot Cluster Generation System using PE Cluster Kit cBot-HS (Illumina) according to the manufacturer’s instructions. After cluster generation, the library preparations were sequenced on an Illumina NovaSeq 6000 Platform (Illumina, San Diego, CA, USA) using a paired-end 150 run (2×150 bases).

### Differential Expression Analysis

Ensembl gene identifiers were converted to gene symbols, and identifiers with duplicated or missing gene symbols were removed from the analysis. Exploratory analyses were performed on the gene-level read count data, and lowly expressed genes were removed from the data set. The EDASeq R package (Risso *et al*, 2011) was used to create a SeqExpressionSet object based on the read counts, then upper quantile normalization was applied. The RUVSeq R package (Risso *et al*., 2011) was then applied using k = 1 and a set of 14 mouse housekeeping genes identified by Ho and Patrizi (Ho & Patrizi, 2021) to identify factors of unwanted variation that were included as a covariate in the differential expression analysis performed with edgeR (McCarthy *et al*, 2012; Robinson *et al*, 2010). Differentially expressed genes were chosen based on a false discovery rate threshold of q < 0.05. R 4.0.5 was used for all analyses (https://www.R-project.org).

### Gene Set Enrichment Analysis (GSEA)

Based on the edgeR output, GSEA software was applied to perform pathway analyses (Mootha *et al*, 2003; Subramanian *et al*, 2005). The genes in the edgeR output were ordered based on a signed version of the likelihood ratio statistic. In brief, the signed likelihood ratio statistic was computed by multiplying the observed likelihood ratio statistic by the sign of the log fold change. Using the KEGG and Hallmark Gene Sets from the Molecular Signatures Database (https://www.gsea-msigdb.org/gsea/msigdb/index.jsp), a GSEA pre-ranked analysis was performed.

### Metabolite Profiling

#### Metabolite extraction

A metabolite extraction was carried out on each sample based on a previously described method(Pacold *et al*, 2016). An approximate cell count (5×10^6^ cells for all samples) of the samples was used to scale the metabolite extraction to a ratio of 1×10^6^ cells / 1 mL extraction solution [80% LCMS grade methanol (aq) with 500nM labelled amino acid internal standard (Cambridge Isotope Laboratories, Inc., Cat No. MSK-A2-1.2)]. The lysis was carried out in two steps as follows. First, 1 mL of freezing extraction solution was added to the tubes containing each pellet. That suspension was transferred to tubes along with zirconium disruption beads (0.5 mm, RPI) and homogenized for 5 min at 4°C in a BeadBlaster™ with a 30 s on / 30 s off pattern. The resulting lysate was then diluted to a fixed concentration of 1×10^6^ cells / 1 mL in a new tube with disruption beads in that same buffer and then re-homogenized in the same way as above. The homogenate was centrifuged at 21,000 x g for 3 min, and 450 μL of the supernatant volume was transferred to a 1.5 mL microfuge tube for speed vacuum concentration, no heating. The dry extracts were resolublized in 50 μL of LCMS grade water, sonicated in a water bath for 2 min, centrifuged as above, and transferred to a glass insert for analysis.

#### LC-MS/MS methodology

Samples were subjected to an LC-MS/MS analysis to detect and quantify known peaks. The LC column was a Millipore™ ZIC-pHILIC (2.1 x150 mm, 5 μm) coupled to a Dionex Ultimate 3000™ system and the column oven temperature was set to 25°C for the gradient elution. A flow rate of 100 μL/min was used with the following buffers; A) 10 mM ammonium carbonate in water, pH 9.0, and B) neat acetonitrile. The gradient profile was as follows; 80-20%B (0-30 min), 20-80%B (30-31 min), 80-80%B (31-42 min). Injection volume was set to 2 μL for all analyses (42 min total run time per injection). MS analyses were carried out by coupling the LC system to a Thermo Q Exactive HF™ mass spectrometer operating in heated electrospray ionization mode (HESI). Method duration was 30 min with a polarity switching data-dependent Top 5 method for both positive and negative modes. Spray voltage for both positive and negative modes was 3.5kV and capillary temperature was set to 320°C with a sheath gas rate of 35, aux gas of 10, and max spray current of 100 μA. The full MS scan for both polarities utilized 120,000 resolution with an AGC target of 3×10^6^ and a maximum IT of 100 ms, and the scan range was from 67-1000 *m*/*z*. Tandem MS spectra for both positive and negative mode used a resolution of 15,000, AGC target of 1×10^5^, maximum IT of 50 ms, isolation window of 0.4 m/z, isolation offset of 0.1 m/z, fixed first mass of 50 m/z, and 3-way multiplexed normalized collision energies (nCE) of 10, 35, 80. The minimum AGC target was 1×10^4^ with an intensity threshold of 2×10^5^. All data were acquired in profile mode.

#### Data analysis

Metabolomics data were processed with an in-house pipeline for statistical analyses and plots were generated using a variety of custom Python code and R libraries including: pheatmap, MetaboAnalystR, manhattanly. Peak height intensities were extracted based on the established accurate mass and retention time for each metabolite as adapted from the Whitehead Institute, and verified with authentic standards and/or high-resolution MS/MS manually curated against the NIST14MS/MS and METLIN spectral libraries. The theoretical m/z of the metabolite molecular ion was used with a ± 10 ppm mass tolerance window, and a ± 0.2 min peak apex retention time tolerance within the expected elution window (1–2 min). To account for sample-to-sample variance in the estimated cell counts, a sum-normalization step was carried out on a per-column (sample) basis. Detected metabolite intensities in a given sample were summed, and a percentage intensity was calculated for each metabolite (custom Rscript). The median mass accuracy vs the theoretical m/z for the library was -0.7 ppm (n = 90 detected metabolites). Median retention time range (time between earliest and latest eluting sample for a given metabolite) was 0.23 min (30 min LCMS method). A signal to noise ratio (S/N) of 3X was used compared to blank controls throughout the sequence to report detection, with a floor of 10,000 (arbitrary units). Labeled amino acid internal standards in each sample were used to assess instrument performance (median CV% = 5%).

### Statistics

All statistical tests were conducted using GraphPad Prism 9 and data presented as mean ± SEM. When comparing two groups the Student’s t-test was used. If greater than two groups were compared, then One-way analysis of variance was used. For all results with normally distributed data, parametric statistical tests were used. When data were not normally distributed, non-parametric statistical tests were used. Data normality was determined using Prism 9.0. Biological replicates were defined as primary cells isolated from an individual mouse from their respective genotype. Statistically significant differences between groups were identified within the figures where *, **, ***, and **** indicating p-values of < 0.05, < 0.01, < 0.001, and <0.0001 respectively.

## Acknowledgements

We thank Dr. Han Chen from The Pennsylvania State University College of Medicine EM facility for assistance with TEM imaging and Dr. Drew Jones and Mr. Leonard Ash from New York University Langone Health Metabolomics Core Resource Laboratory for assistance with metabolomics experiments. We are grateful to the Flow Cytometry and Informatic and Data Analysis Core facilities from The Pennsylvania State University College of Medicine. This work was supported by National Institutes of Health (MIH) grants R35-HL150778 (to M.T.) and in part by R35-HL161177 (to A.C.S), R01-AI162971 (to ZSMR) and R01-AI097302 and R01-AI130143 (to SF).

## Material Availability Statement

All unique material generated in the study, including Orai3^fl/fl^ mice, CRISPR-Cas9 Orai knockout A20 clones and other reagents are available from the lead author upon request.

## Figure Legends

**Figure 3- Figure Supplement 1.**
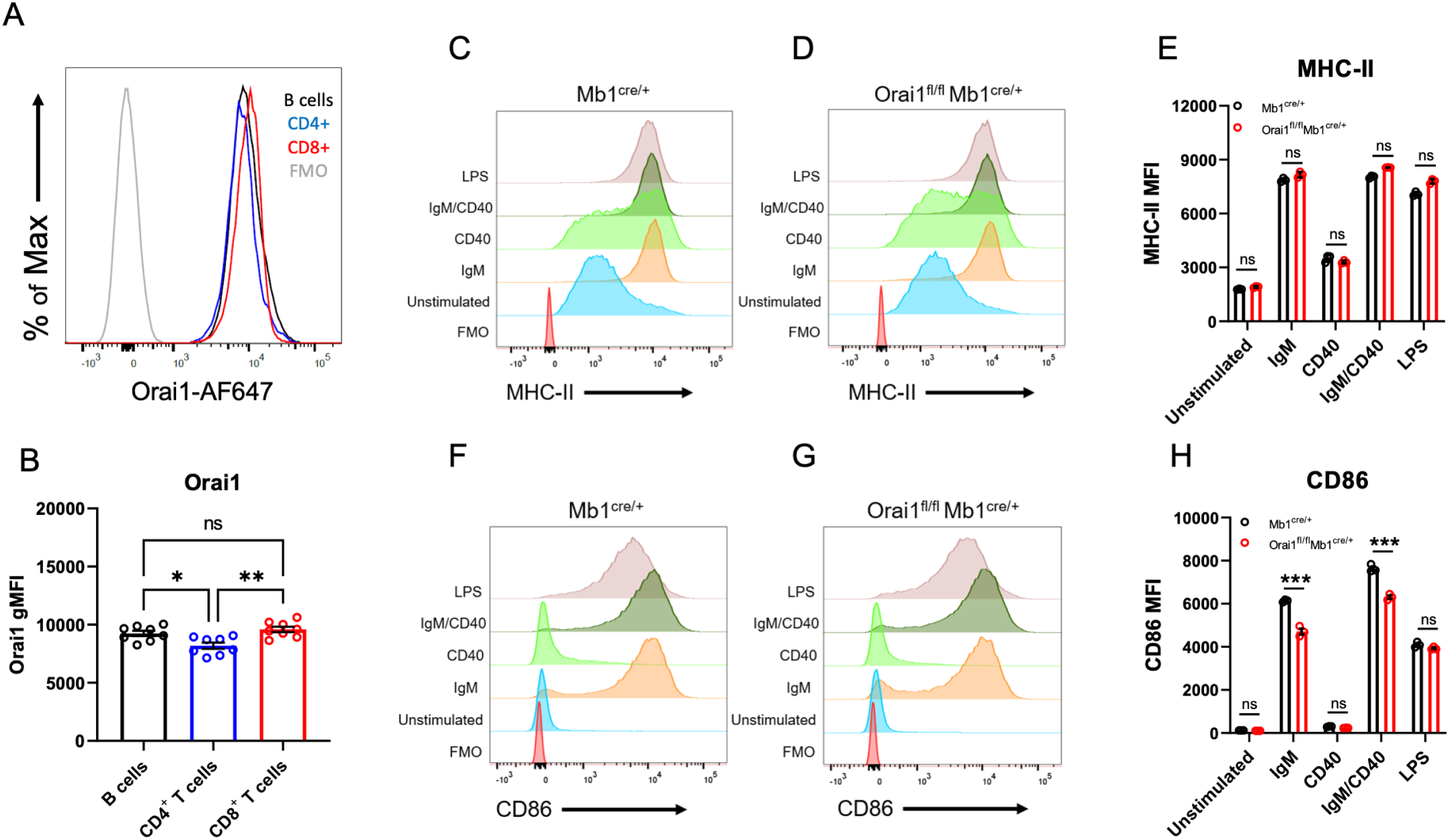
Loss of Orai1 does not overtly affect B cell activation. (A) Representative histograms of Orai1 fluorescence gated on B220^+^ B cells, CD3^+^CD4^+^ T cells, and CD3^+^CD8^+^ T cells from *Mb1*^*cre/+*^ mice. (B) Quantification of Orai1 mean fluorescence intensity (MFI) from each condition shown in (A) (n = 8 biological replicates for each; one-way ANOVA). (C-D) Flow cytometry histograms of MHC-II expression on isolated B cells from (C) *Mb1*^*cre/+*^ and (D) *Orai1*^*fl/fl*^ *Mb1*^*cre/+*^ mice following 24-hour stimulation. (E) Quantification of MHC-II MFI from (C-D) (n = 3 biological replicates for each; Mann-Whitney test). (F-G) Histograms of CD86 expression following 24-hour stimulation. (H) Quantification of CD86 MFI from (F-G) (n = 3 biological replicates for each; Mann-Whitney test). All scatter plots are presented as mean ± SEM. For all figures, *p<0.05; **p<0.01; ***p<0.001; ns, not significant.

**Figure 6- Figure Supplement 1.**
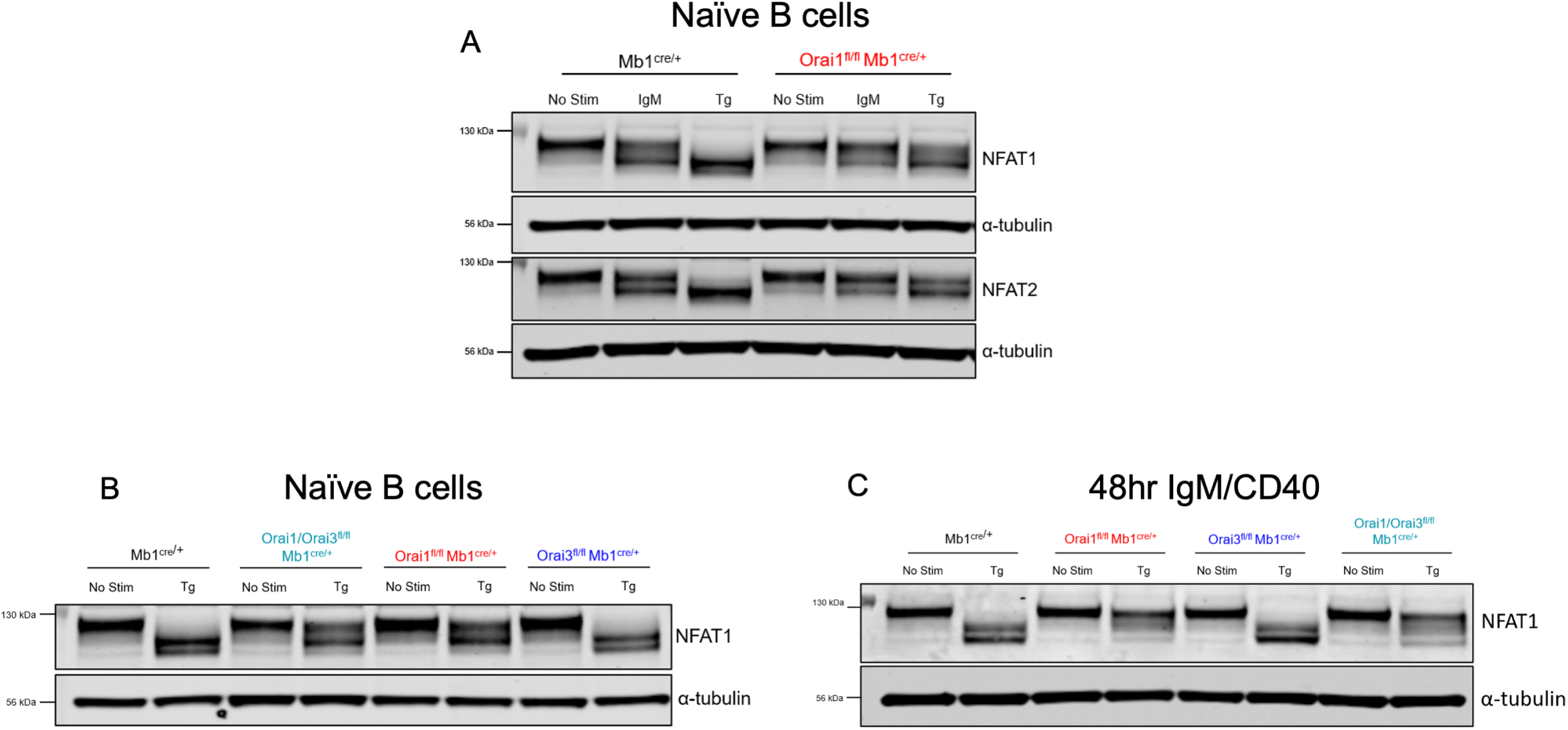
NFAT activation in primary Orai1/Orai3-deficient B cells. (A) Western blot analysis of NFAT1, NFAT2, and α-tubulin in naïve B cells isolated from *Mb1*^*cre/+*^ and *Orai1*^*fl/fl*^ *Mb1*^*cre/+*^ mice. B cells were either unstimulated or treated with 20μg/mL anti-IgM or 2µM thapsigargin (Tg) for 15 minutes before harvesting. (B) Western blot analysis of NFAT1 in naïve B cells isolated from *Mb1*^*cre/+*^, *Orai1*^*fl/fl*^ *Mb1*^*cre/+*^, *Orai3*^*fl/fl*^ *Mb1*^*cre/+*,^ and *Orai1/Orai31*^*fl/fl*^ *Mb1*^*cre/+*^ mice following thapsigargin treatment. (C) Same as in (B) but with B cells stimulated for 48 hours with anti-IgM + anti-CD40.

**Figure 7- Figure Supplement 1.**
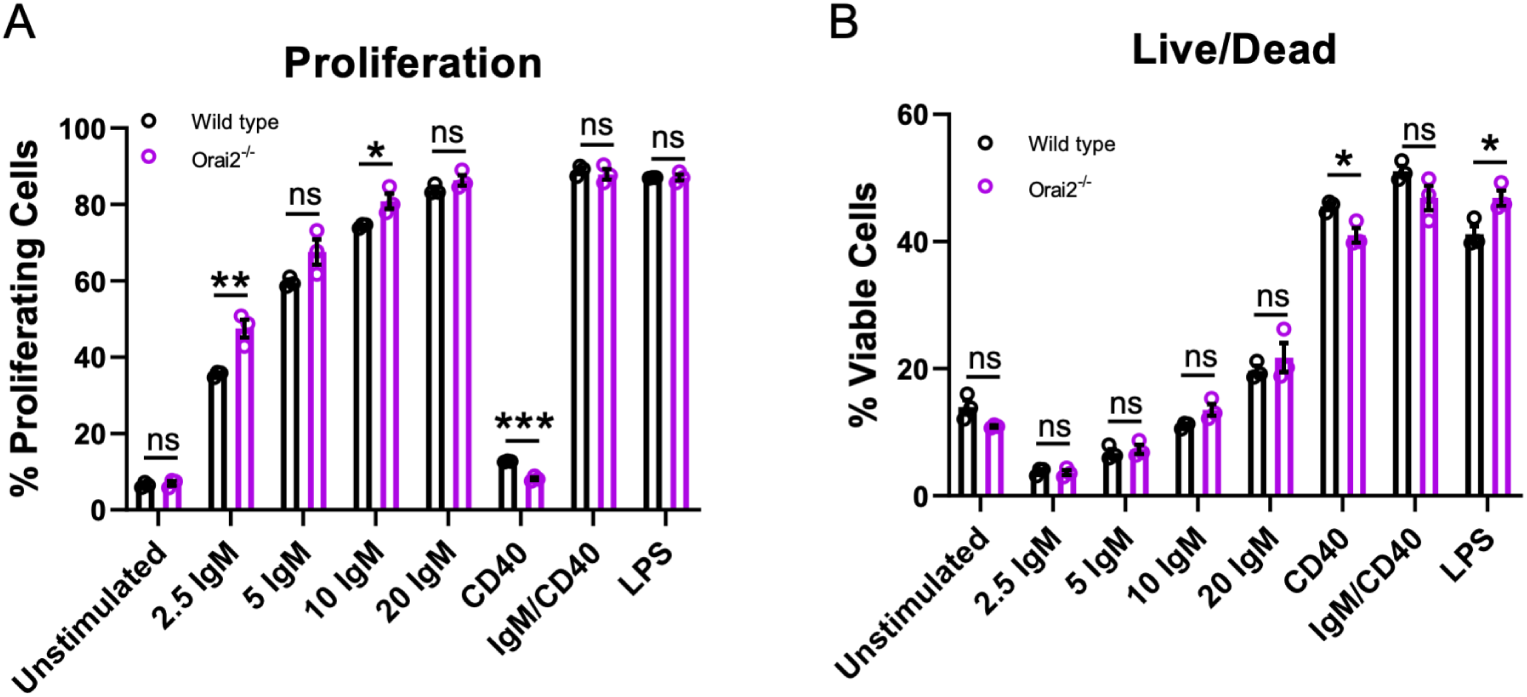
Loss of Orai2 does not alter primary B cell proliferation or viability. (A) Quantification of B cell proliferation by tracking CFSE dilution. B lymphocytes from wild-type and *Orai2*^*-/-*^ mice were loaded with CFSE (3 μM) and stimulated with a titration of anti-IgM antibodies, anti-CD40 (10μg/mL), anti-IgM + anti-CD40, or LPS (10μg/mL). CFSE dilution was determined 72 hours after stimulation for all conditions (n = 3 biological replicates for each; unpaired T-test). (B) Quantification of the percentage of viable cells for each condition in (A) as determined by a Live/Dead viability dye (n = 3 biological replicates for each; unpaired T-test). All scatter plots are presented as mean ± SEM. For all figures, *p<0.05; **p<0.01; ***p<0.001; ns, not significant.

**Figure 8- Figure Supplement 1.**
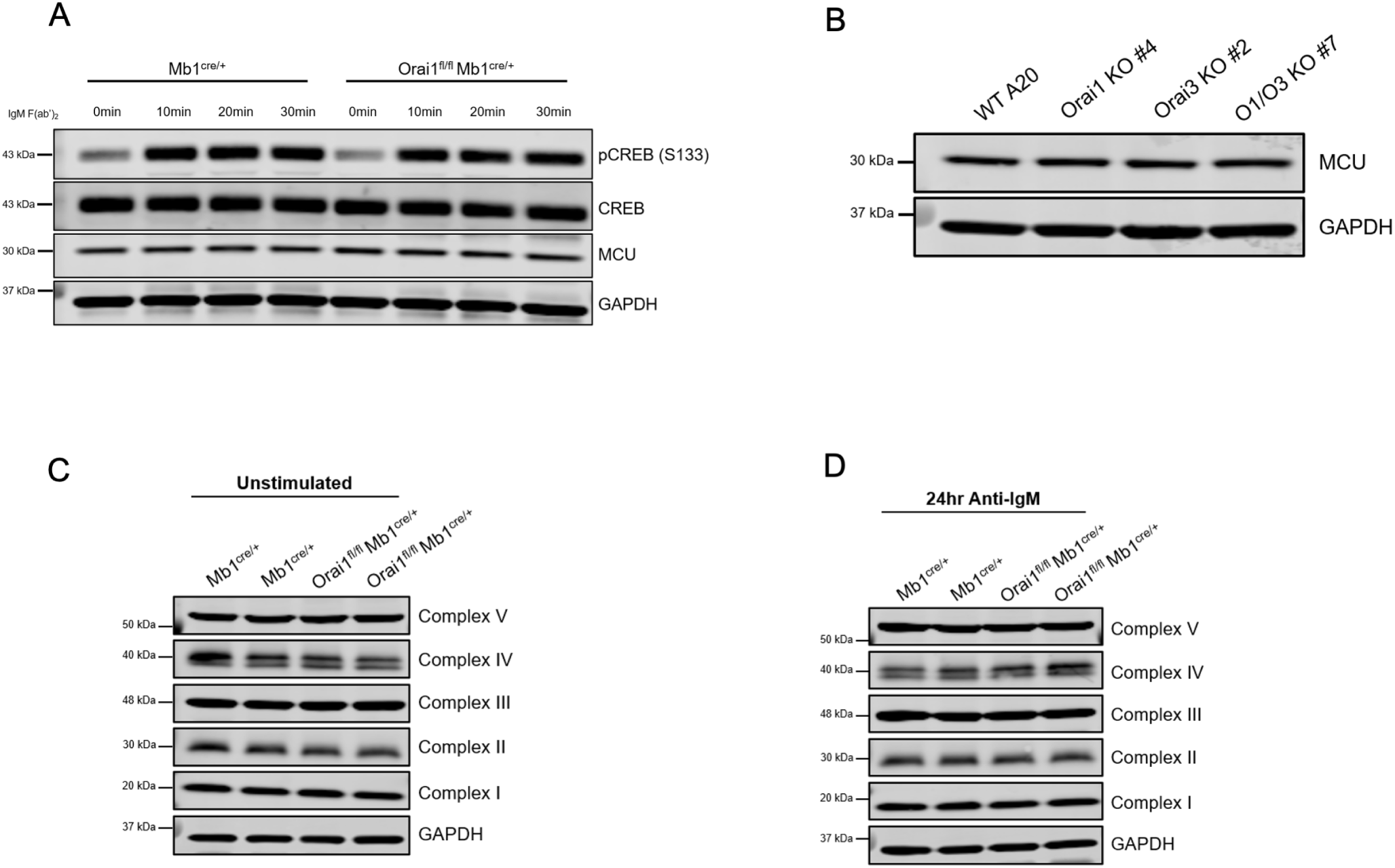
Loss of Orai1 does not alter CREB phosphorylation, MCU expression or expression of electron transport chain proteins. (A) Western blot analysis of total and phosphorylated CREB (S133), MCU, and GAPDH in isolated B cells from *Mb1*^*cre/+*^ and *Orai1*^*fl/fl*^ *Mb1*^*cre/+*^ mice. B cells were either unstimulated or stimulated with anti-IgM (20μg/mL) for 10, 20, or 30 minutes before harvesting. (B) Western blot analysis of MCU and GAPDH protein in single Orai1, and Orai3 knockout and double Orai1/Orai3 knockout A20 cell clones. (C, D) Western blot analysis of electron transport chain components in B cells from *Mb1*^*cre/+*^ and *Orai1*^*fl/fl*^ *Mb1*^*cre/+*^ mice either (C) unstimulated or (D) stimulated with anti-IgM for 24 hours.

**Figure 9- Figure Supplement 1.**
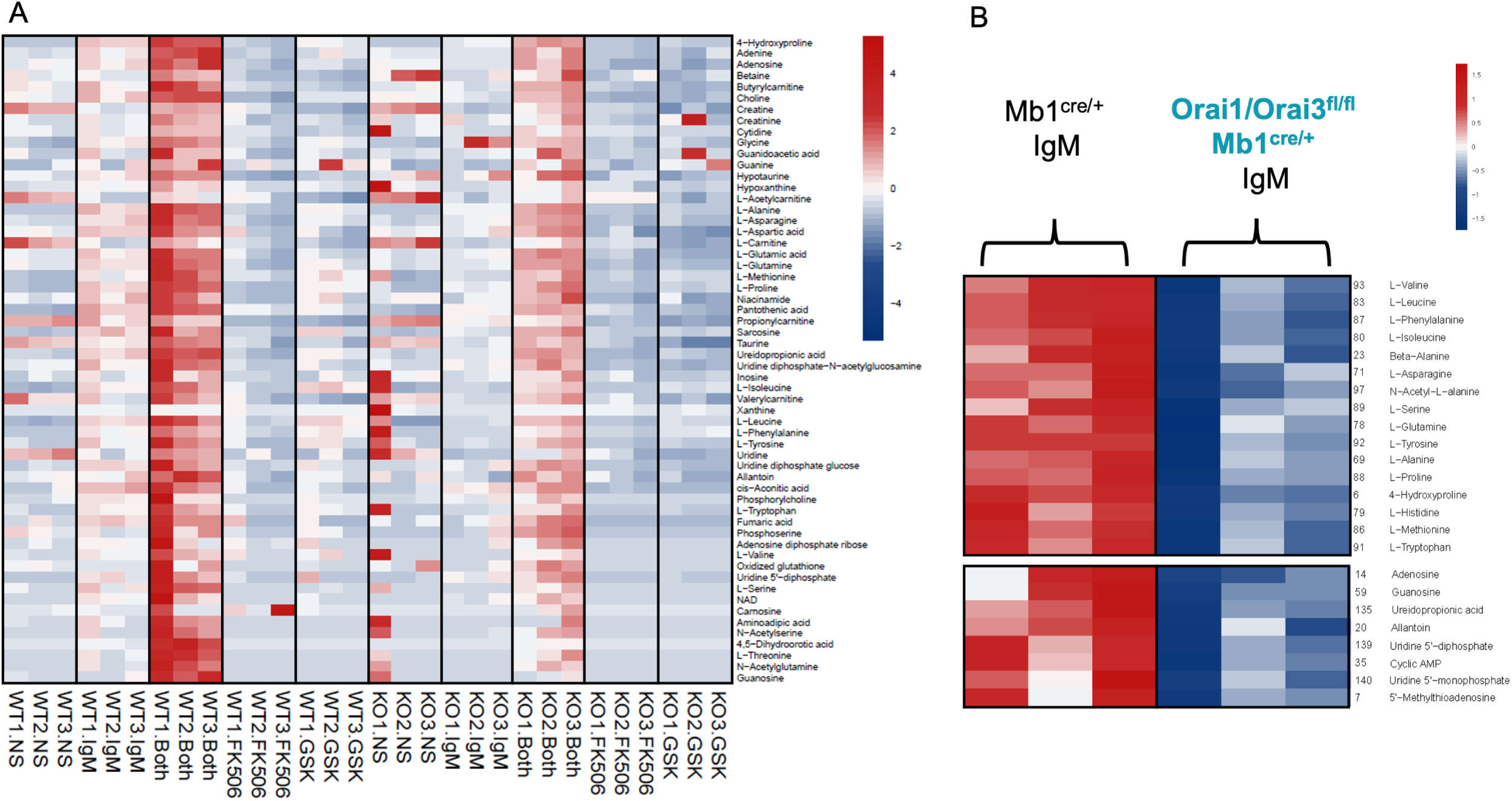
Polar metabolite analysis in Orai1/Orai3 deficient-B cells. (A) Analysis of polar metabolites in B cells from *Mb1*^*cre/+*^ and *Orai1/Orai31*^*fl/fl*^ *Mb1*^*cre/+*^. B cells were either unstimulated or stimulated for 24 hours with anti-IgM, anti-IgM + anti-CD40, anti-IgM + 1µM FK506, or anti-IgM + 10µM GSK-7975A. Profiling was performed on B cells from 3 mice per group. (B) Heat maps of statistically significant amino acids and nucleotide precursors in B cells from *Mb1*^*cre/+*^ and *Orai1/Orai31*^*fl/fl*^ *Mb1*^*cre/+*^ mice following anti-IgM stimulation for 24 hours.

**Source data 1:** Zipped folder containing raw unedited source data of images of all gels and Westerns. Each image is labeled and associated to its corresponding figure in the manuscript. The uncropped images of gels with relevant bands labeled are provided as Excel files and labeled accordingly.

